# SwarmMAP: Swarm Learning for Decentralized Cell Type Annotation in Single Cell Sequencing Data

**DOI:** 10.1101/2025.01.13.632775

**Authors:** Oliver Lester Saldanha, Vivien Goepp, Kevin Pfeiffer, Hyojin Kim, Jie Fu Zhu, Rafael Kramann, Sikander Hayat, Jakob Nikolas Kather

## Abstract

Rapid technological advancements have made it possible to generate single-cell data at a large scale. Several laboratories around the world can now generate single-cell transcriptomic data from different tissues. Unsupervised clustering, followed by annotation of the cell type of the identified clusters, is a crucial step in single-cell analyses. However, there is no consensus on the marker genes to use for annotation, and celltype annotation is currently mostly done by manual inspection of marker genes, which is irreproducible, and poorly scalable. Additionally, patient-privacy is also a critical issue with human datasets. There is a critical need to standardize and automate celltype annotation across datasets in a privacy-preserving manner. Here, we developed SwarmMAP that uses Swarm Learning to train machine learning models for cell-type classification based on single-cell sequencing data in a decentralized way. SwarmMAP does not require any exchange of raw data between data centers. SwarmMAP has a F1-score of 0.93, 0.98, and 0.88 for cell type classification in human heart, lung, and breast datasets, respectively. Swarm Learning-based models yield an average performance of **0.907** which is on par with the performance achieved by models trained on centralized data (*p*-val=**0.937**, Mann-Whitney *U* Test). We also find that increasing the number of datasets increases cell-type prediction accuracy and enables handling higher cell-type diversity. Together, these findings demonstrate that Swarm Learning is a viable approach to automate cell-type annotation. SwarmMAP is available at https://github.com/hayatlab/SwarmMAP.

## 1 Introduction

Recent technological advances in single-cell sequencing have led to a plethora of scientific discoveries improving our understanding of human tissue and diseases [1], including COVID [2, 3], lung [4], cardiovascular [5–7], renal diseases [8], and cancer [9–11] at singlecell resolution. Typical single-cell analysis pipelines use unsupervised clustering, followed by cell-type annotation of identified clusters based on the expression level and specificity of selected marker genes [12]. Cell-type annotation is still primarily a manual effort in which subject experts review marker genes per cluster to annotate cell types. There is no consensus on marker genes and their importance for cell type annotation yet. This reduces reproducibility as the selection of marker genes and their importance for annotation is dependent on the expert annotating the data and can vary from person to person. Furthermore, with increasing amounts of data emerging from different labs, this approach is not scalable or transferable. Additionally, secure data sharing and maintaining data privacy is a critical issue while working with human patient data [13].

To leverage the full potential of multiple studies, tools to generate standardized celltype annotation while maintaining data privacy are needed to unify and compare data across studies. Furthermore, to increase reproducibility, scalability, and limit individual bias, manual annotation of cell clusters should be increasingly replaced by machine learning models that automatically assign individual cells to a cell type [14–17]. Some tools have also been developed to map new unannotated data to a reference data set [18–20]. However, it is challenging to train a universal machine learning model to classify cell types based on individual single-cell sequencing datasets due to the underlying technical batch effects. Moreover, generalizable machine learning models need to be trained on large, multi-centric, and diverse datasets to account for this variability. The usual procedure to create such datasets is centralized data collection. This requires multiple participating institutions to send their data to a single location. Such data transfer can create practical, legal, and even ethical problems and is often a rate-limiting step to train machine learning models in biology and medicine [21].

Swarm learning is a computational technique to co-train machine learning models at multiple institutions in a decentralized way, without exchanging underlying data[22]. Swarm learning does not require a central coordinator of the network and thus avoids monopolization of resources and machine learning models [23]. In medical image and computational pathology analysis, Swarm learning has been shown to enable a high performance of machine learning models, which is on par with models trained in a centralized way [23–25]. Ultimately, Swarm learning could enable training of machine learning models in a massively parallelized way, increasing the resilience of the training process and democratizing access to the resulting models.

Here, we show that Swarm learning can be efficiently applied to train machine learning models for cell type classification based on single cell sequencing data. We evaluate this on human data from multiple organs generated by different research centers. Our tool, SwarmMAP, shows high accuracy for cell-type classification in a privacy-preserved setting where patient data is not shared among users. SwarmMAP enables comparative analyses across datasets, enabling novel discoveries in single-cell datasets while maintaining patient privacy.

## 2 Materials and Methods

### 2.1 Overview of the workflow

SwarmMAP is a Swarm learning based method to classify cell-types in single-cell transcriptomic data. It is trained in a supervised learning manner for each organ. We access the utility of SwarmMAP in both local learning (LL) and Swarm learning (SL) settings using the same data pre-processing pipeline, and classifier is used to compare performance (Figure 1A). In LL, a single model is trained using a common training dataset (Figure 1B) while in SL, each agent keeps its data and model private (Figure 1C), as a blockchain-enabled Swarm learning framework allows each agent’s model to learn from the other models. To assess model performance, the predicted annotation is compared with the annotation provided by the authors in the test dataset in a study-specific manner.

**Fig. 1:**
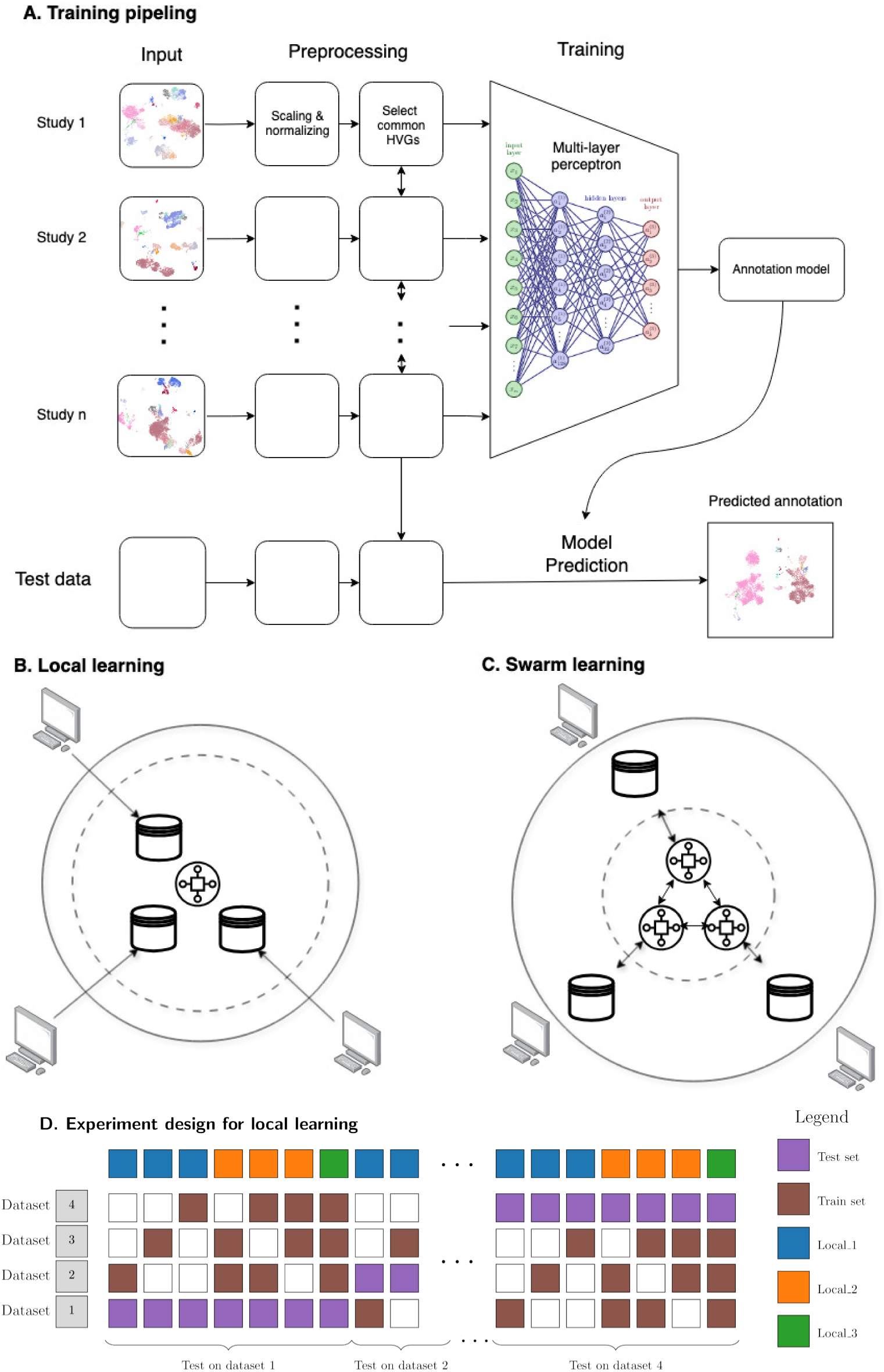
Overview of the SwarmMAP workflow. (A) Pipeline for training our annotation classifier: scaling and normalization is done independently for each study, except for selecting the highly variable genes (HVGs), where the union of the top 2000 HVGs across studies is used; (B) Local learning (LL) framework, where all the data is shared to learn a single model (dashed inner circle); (C) Swarm learning (SL) framework, where no data is shared and decentralized sharing of models’ parameters is enable by a blockchain (dashed inner circle); (D) Experimental design for local learning (LL): for each organ, all combinations of train and test sets are used. The settings where 1, 2, or 3 training sets are used are termed Local_1, Local_2, and Local_3. For Swarm leaning (SL), all 3 training sets are used in a data privacy-preserving manner.

### 2.2 Dataset description

SwarmMAP is trained and tested on single-cell transcriptomic data from human heart, lung, and breast (Supplementary Table 1) totaling 284 donors and 1,956,243 cells. Each collection consists of 4 separate studies. Lung and breast collections are taken from the human lung cell atlas [4] and breast atlas [9], respectively. The heart collection is created from individual datasets, where the cell-types provided by individual studies have been standardized. Four studies were selected from each collection, with varying sample sizes and cell composition (S12). Training and testing is performed at two annotation levels: cell types and cell subtypes, which are used as the ground truth for the classifier. The datasets and their cell-types are represented in Supplementary Figures S1 and S2 using UMAP [26].

### 2.3 Preparation of labels from cell type annotation

We perform the classification of cell labels at two annotation levels: a coarse annotation level, “cell types”, and a finer annotation level, “cell subtypes”. Each annotation level is trained independently using its own model.

All datasets included in this study come with their own annotations by the respective authors. For the heart atlas, the level 1 annotation (“Annotation_1”) was used as cell types (14 cell types). The level 1 annotation (“Subclustering”) contained 65 cell subtypes, which are too specific for cell type annotation. Thus, the subtypes were grouped together to arrive at 17 subtypes of cells. The “Subclustering” labels were manually matched to their closest cell ontology terms [27]. Then, hierarchical clustering was performed across the 4 studies to measure distance between subtypes and the closest subtypes are merged together, the names being set to their respective common ancestors in the cell ontology. For heart, we obtained 17 cell subtypes from 65 initial “Subclustering” labels.

For lung and breast atlases, the same merging process was used, except that cell ontology terms were already available. For the lung atlas, the initial “cell_type” label with 50 categories was gradually merged to obtain 24 cell subtypes, and then further to obtain 17 cell types. For the breast atlas, the initial “cell_type” label with 26 categories was gradually merged to obtain 22 cell subtypes and 14 cell types.

The resulting annotations have the number of classes comparable between the heart, lung, and breast collections: 14, 17, and 14, respectively, for cell types, and 17, 24, and 22, respectively, for cell subtypes. The hierarchical clustering of the final annotations are provided in dendrograms (Supplementary Figure S7) and the correspondence between cell types and cell subtypes is represented in a Sankey diagram (Supplementary Figure S5).

### 2.4 Data preprocessing for supervised learning

For each data collection, we apply the following preprocessing steps before training.

- **Suspension type**: Suspension types (single-cell or single-nuclei) are filtered to ensure that all observations in every data collection have the same suspension type. The heart collection consists of single nuclei, and the lung and breast collections consist of single cell data.
- **Tissue**: Data are also filtered with respect to the tissue. For the heart, only cells from the left ventricle are selected. For the lung, only cells from the lung parenchyma are selected. The breast collection has no tissue specification.
- **Quality Control**: Standard quality control (QC) of counts is performed and cells deemed outliers are filtered out. Four QC metrics are used: log1p value of total counts, log1p value of number of genes by counts, percentage of counts in the top 20 genes, and percentage of reads mapped to mitochondrial genes. For each metric, each cell whose metric is smaller or greater than a margin of five times the median absolute deviation around the median value is set as an outlier. Furthermore, all cells with more than 8 percent of reads mapped to mitochondrial genes are also set as outliers.
- **Feature selection**: For each study, the 2000 most highly variable genes (HVGs) are computed. The choice of 2000 features constitutes a trade-off between model simplicity and model complexity. The effect of the number of HVGs is reported in Supplementary Figure S9. The set of features is then defined as the union of these genes in all studies (3516, 3566, and 3174 features for heart, lung, and breast, respectively). Consequently, in the SL setting, each model is trained after sharing the union of all other agents’ HVGs. We consider that sharing this information constitutes no privacy breach concerning the expression data (see section 4). Moreover, the initial number of HVGs used can have an effect on the model performance and is the result of a trade-off between model complexity and generalizability. Supplementary Figure S9 displays the F1 scores for each cell type for several number of HVGs: 200, 500, 1000, and 2000. For the heart and lung, performance increases continually as the number of HVGs increases, but with diminishing returns. For breast collection, interestingly, the number of HVGs showed no effect on performance. Overall, these results suggest that a higher number of HVGs, is beneficial for classification. However, this comes at the cost of computing time and model complexity, we choose the standard value of 2000 HVGs throughout [28].

The final dataset consists of 1,196,647 (1,117,502), 243,031, 516,565 cells in heart (heart subtype), lung and breast collections, respectively (see S12) from 144, 51, and 89 donors (see Supplementary Table 2), respectively. Finally, raw counts are normalized (10,000 counts per cell) and scaled using the log(1+x) transform.

### 2.5 Experimental design

The experiment design for local learning (LL) is detailed in Figure 1D. Each column represents an experiment and the experiment design (28 experiments in total) and is applied to each organ separately. Each experiment only uses one dataset as a test set. Then, for each choice of test set, all combinations of training sets are used, that is, training on 1 (“Local_1”, 12 experiments), 2 (“Local_2”, 12 experiments), and 3 (“Local_3”, 4 experiments) datasets. This design compares the classification performance as a function of the number of cells, averaging out the difference in cell count between datasets. For Swarm learning (SL), only the four experiments designs with 3 training sets are used, using all combinations of test set.

### 2.6 Machine learning classifier model

Briefly, the classifier is trained in both the local and Swarm learning setups separately. In the local learning (LL) setup, classification performance is evaluated when training on 1, 2, or 3 datasets, where the fourth dataset is used for testing. These simulation settings are called Local_1, Local_2, and Local_3, respectively. All possible combinations of training and testing datasets are used. In the Swarm learning (SL) setup, 3 datasets are used for training and one for testing. For each combination of train and test datasets, the validation set is obtained by splitting the training data into train and validation sets.

The model used for classification is a multi-layer perceptron (MLP) classifier. We use two fully connected inner layers with 128 and 32 neurons, respectively. We use a tanh activation function and a cross-entropy loss. Optimization is performed using the Adam optimizer. After hyperparameter fine-tuning using cross-validation on the heart dataset with cell types as labels, the following parameters are used: a learning rate of 1e-3 and no weight decay; a batch size of 128; 100 training epochs. The following alternative configurations were tested:

- using dropout for regularization (dropout rates of 0.25 and 0.5);
- using weight decay with the AdamW optimizer (value between 1e-3 and 1e-7);
- using weighted class resampling to counterweight the class imbalance; and
- including inverse class proportions as weights in the loss function;
- using different batch sizes (32, 64, or 256).

These did not produce a significant improvement in classification.

### 2.7 Swarm Learning

Swarm learning (SL) allows decentralized collaborative training of machine learning (ML) models on multiple physically distinct computing systems (peers) [29]. Here, we implemented SL using three separate peers, representing the institutions. Each peer independently trained a machine learning model on its proprietary dataset, with no raw data shared between peers. During training, model weights and biases were exchanged in multiple synchronization events (sync events). These sync events occurred at the end of each synchronization interval, defined as a fixed number of training batches. At each sync event, the model weights were averaged, and training was resumed at each peer using the updated parameters. To account for differences in sizes of the datasets, we applied a weighted SL approach, where the contributions of each peer were scaled by a weighting factor proportional to the size of its dataset. Motivated by previous studies on pathological and radiology data [23, 30]. This approach ensured balanced contributions from peers with varying dataset sizes, where larger datasets are not overrepresented in the final model. After completing all training epochs, a final round of model merging is performed, providing all peers with a unified model. The Hewlett Packard Enterprise (HPE) SL framework was utilized, which consists of five major components: the ML node, the SL node, the Swarm Network (SN) node, identity management, and HPE license management. The ML node defines the ML model and access to the data. Where the SL node process handles the parameter sharing, while the SN node process manages peer communication. To manage global model state information and enable decentralized parameter merging, an Ethereum blockchain (https://ethereum.org) was employed. Unlike traditional federated learning, SL does not rely on a central server; instead, smart contracts facilitate the selection of peers for parameter merging. All processes were executed in Docker containers. A detailed description of this process and instructions for reproduction can be found under https://github.com/KatherLab/swarm-learning-hpe/tree/dev_single_cell.

### 2.8 Comparison with state-of-the-art cell type classifiers

The Swarm learning framework can be applied using any machine learning model, the only constraint being that no data are shared between agents. Thus, SwarmMAP can be built using a variety of classifiers. There have been many approaches to cell type classification in single-cell RNA sequencing data. Garnett [31] uses marker genes already curated to annotate cells using a regularized multinomial linear classifier. ACTINN [32] uses an MLP classifier with 3 hidden layers (100, 50 and 25 neurons, respectively). Celltypist [14] uses L2 regularized logistic regression. Supervised Contrastive Learning for Single Cell (SCLSC) [33] employs contrastive learning to learn an embedding representation for cell types and a KNN classifier to annotate cells. devCellPy is a machine learning-enabled pipeline for automated annotation of complex multilayered single-cell transcriptomic data, based on the XGBoost classifier. scTab [34] introduces a feature-attention-based classifier model for single cell transcriptomic data based on TabNet, a deep learning classifier for tabular data [35]. The classification performance of TabNet was compared with several models, including XGBoost [36] and MLP. [34] found that the feature attention-based classifier model outperformed the other models in the context of large-scale and curated datasets (significant differences in the macro F1 score), but with minor differences in F1 values (0.83 for scTab, 0.81 for XGBoost, 0.80 for MLP).

However, scTab is trained on very large datasets with between 10^3^ and 10^6^ cells per cell type, while SwarmMAP is considering the more challenging setting where some cell types have cell counts of the order of 10^2^, and 10^1^ for some cell types. Since in general settings, XGBoost is preferred over TabNet [37] and is considered to be more flexible for classification task, we chose to compare the performance of our MLP classifier with XGBoost. Specifically, XGBClassifier classifier was used from the XGBoost Python package, with default parameters. XGBoost was compared to MLP for cell type classification in the LL framework, on the same data (3 organs, see Section 2.4 and with the same experiment design (Local_1, Local_2, and Local_3, see Section 2.5). MLP compares slightly favorably to XGBoost while being faster to train by a factor of 2 to 4 (see Supplementary Figure S3). Thus, SwarmMAP is built using an MLP classifier.

### 2.9 Data and code availability

The 12 datasets are publicly available from CellxGene^1^ and the Broad Institute Single Cell portal^2^. The download links are provided in Supplementary Table 3. The processed datasets will be made available on Zenodo upon publication of the manuscript. The SwarmMAP method and the code to reproduce the results in this study are available at https://github.com/hayatlab/SwarmMAP.

## 3 Results

### 3.1 Machine learning-based cell type prediction in multiple datasets

The average weighted F1 score for main cell-type classification in heart datasets are 0.947, 0.957, and 0.958 when training on 1, 2 or 3 (Local_3) datasets, respectively. The corresponding values for cell subtype classification are 0.961, 0.968, and 0.972 respectively (Figure 2). Similar values are obtained from the lung and breast datasets (Figure 2). Here, weighted F1 score (which averages the F1 score for each class weighted by the support of the class) was used as the main classification metric. Results from micro F1 score (based on global true positives, false negatives, and false positives) and the macro F1 score (unweighted average of the F1 score for each class) are also provided in Supplementary Figure S4.

**Fig. 2:**
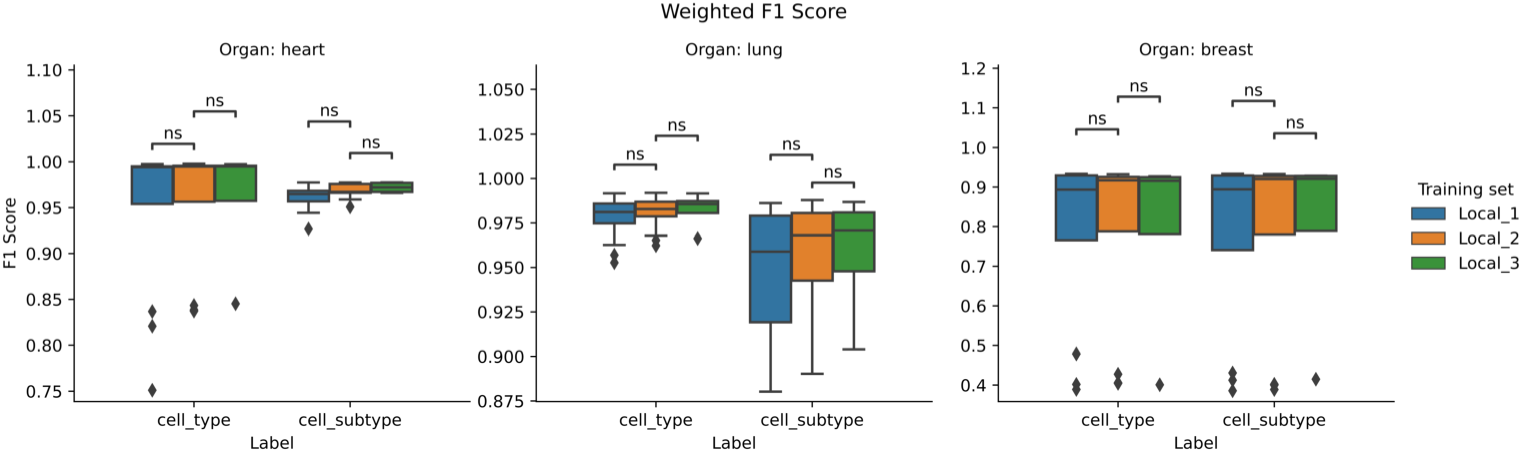
Classification performance of the local learning framework. Weighted F1 scores are averaged over all simulation runs. F1 score distributions for classifying main cell types and subtypes using Local_1 (training on 1 dataset), Local_2 (training on 2 datasets), and Local_3 (training on 3 datasets) setting in local learning.

Mann–Whitney *U* tests were performed to compare F1 scores between the different simulation settings. Despite the increase in mean performance as more datasets are used for training, the differences are not statistically significant, owing to the small sample size of the simulation runs (12, 12, and 4 respectively). Moreover, some cell types are hard to classify (see Section 3.1.2), making the distribution of F1 scores more dispersed. This is especially true for breast data, in which classification is harder (see Section 3.2.3). However, there is an increase in the averaged F1 scores, especially for heart subtypes, lung main cell types, and subtypes. The respective average scores of Local_1, Local_2, and Local_3 are 0.961, 0.968, and 0.972 for heart subtypes; 0.978, 0.981, and 0.982 for lung types; and 0.945, 0.954, and 0.958 for lung subtypes (see Supplementary Table 4 for weighted F1 scores).

#### 3.1.1 Classification performance increases as more datasets are used for training

#### 3.1.2 Classification performances vary greatly between cell-types

Model performance (F1-score) per cell type varies between different cell types (Figure 3). The corresponding results for cell subtypes are represented in Supplementary Figure S11. In particular, for heart data, “Epicardium” cells and “Ischemic cells (Myocardial infarction)” are difficult to classify, in line with the scarcity of these classes, their uneven distribution among datasets (see cell count barplots in Figure S8), and the difficulty to define ischemic cells biologically (see UMAP representation in Figure S1). In addition, the ischemic cells is a collection of cells from different lineages including cardiomyocytes, epithelial, etc. Thus, they do not have well-defined marker genes and are thus inherently difficult to classify. For the lung data, all cell types are well classified, except “respiratory basal cells”, which are classified as “epithelial cells”. For the breast data, classification is more challenging, with four cell types with F1 score below 0.5: mature alpha-beta T cells, mature B cells, naive thymus-derived CD4-positive, alpha-beta T cells, and capillary endothelial cells. Some other cell types like endothelial tip cells and macrophages have a high disparity in classification performance between simulation runs (see Section 3.2.3).

**Fig. 3:**
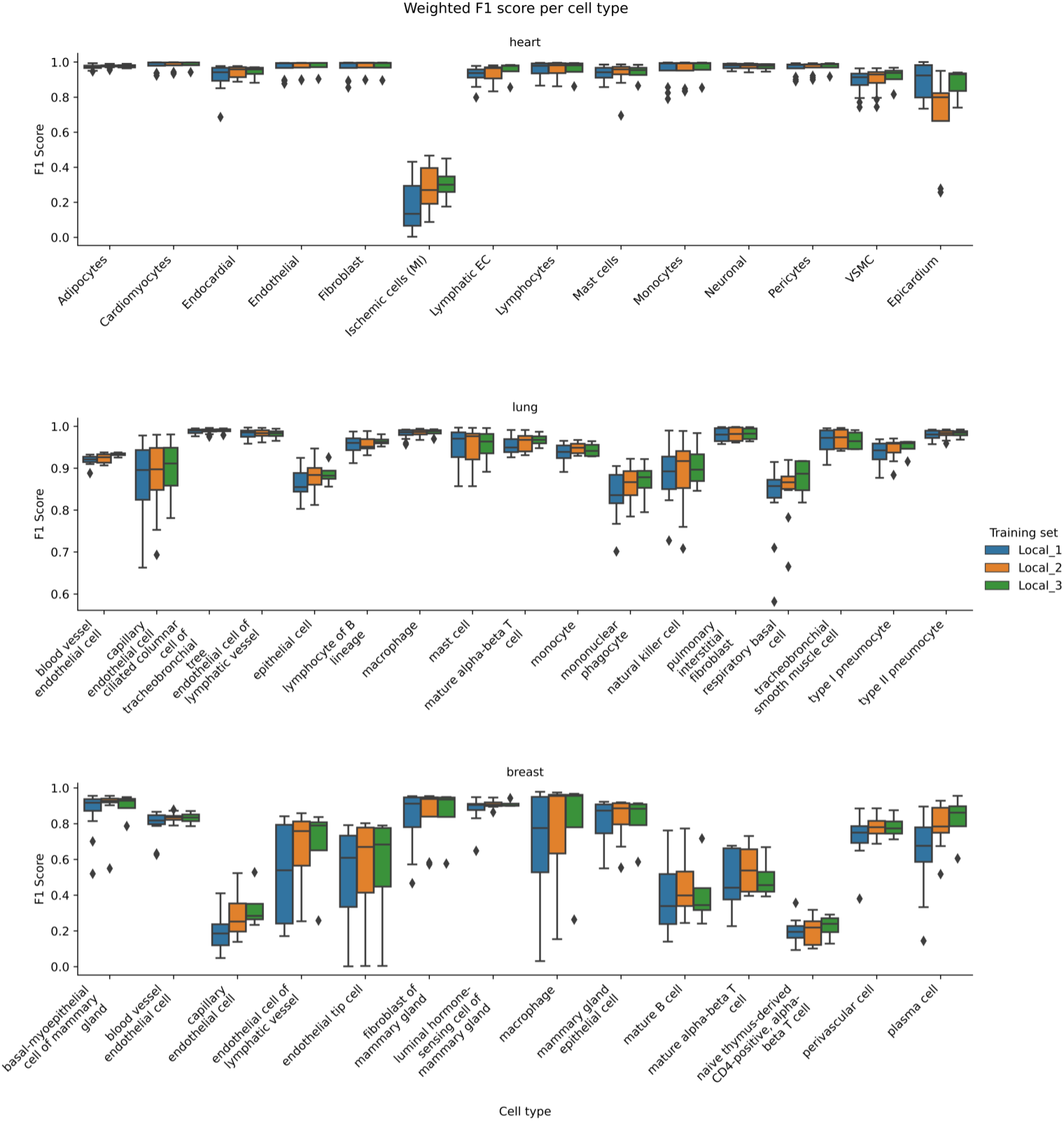
Classification performance of the local learning framework. F1 scores are averaged over all simulation runs for each cell type. Overall, there is a notable disparity in classification accuracy between cell types.

#### 3.1.3 Cell type prediction performance improves as cell count increases

To investigate that increasing the sample size of a cell type improves its classification accuracy, we evaluate the link between classification performance and the rarity of cell types. For each cell type and averaged over all studies, we compute the Gini impurity index, which is a measure of the rarity of the cell type (higher values are rarer classes). Then we computed the F1 score for each cell type and each study, averaged over all simulation runs (28 runs, combining Local_1, Local_2, and Local_3 together). Figure 4 represents the F1 score as a function of the Gini impurity index of cell types. The results show that the classification performance increases as the rarity of the cell types increases.

**Fig. 4:**
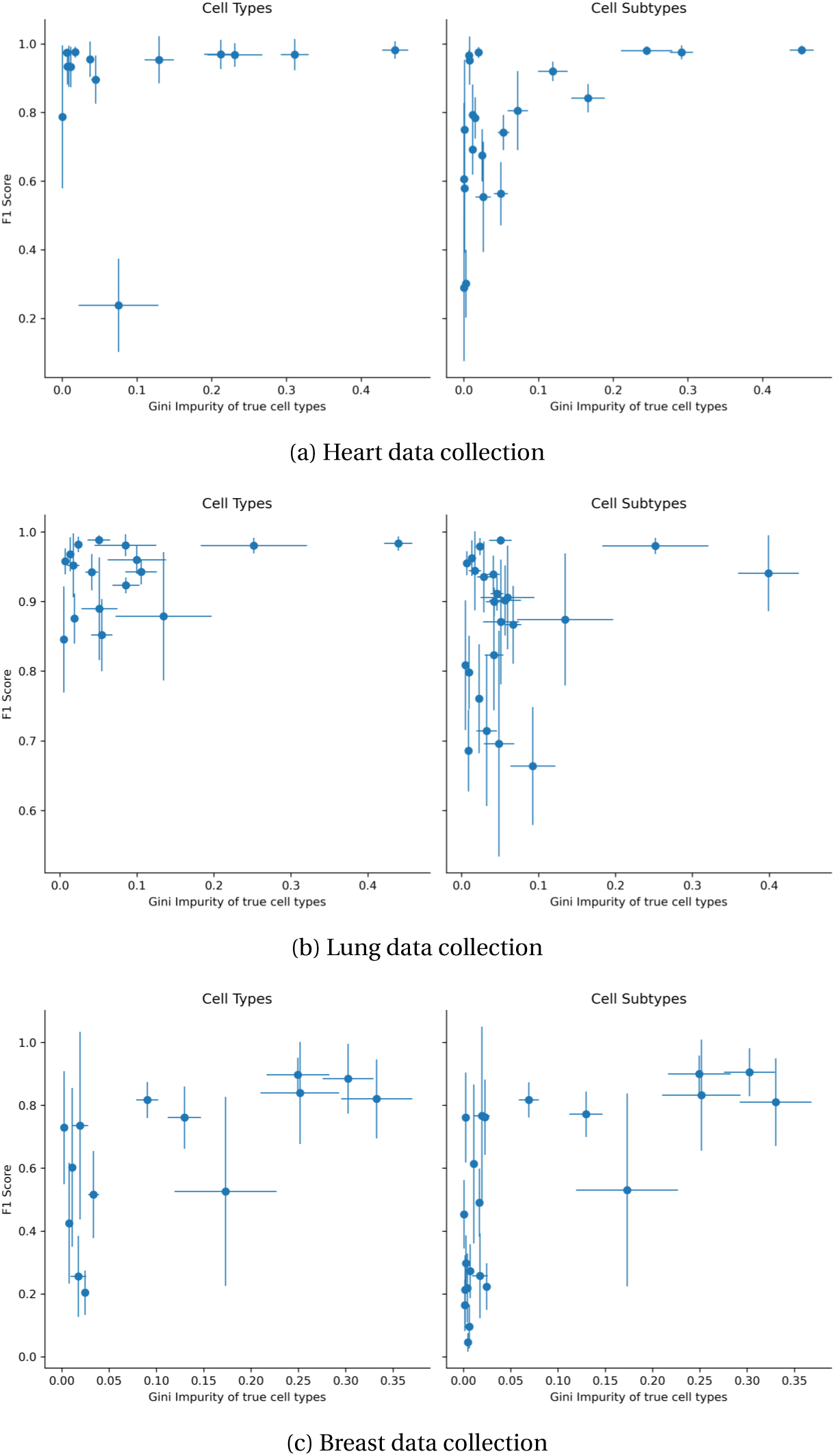
Classification performance as a function of the rarity of the cell types. The x-axis represents the Gini impurity index of true cell types (higher values are rarer classes). The y-axis represents the F1 score. Each dot is a cell type and the values are averaged over all studies for the Gini indices and all simulation runs (Local_1, Local_2, Local_3) for the F1 scores. 13

To quantify this, a linear model was fitted to the data for each organ and level (considering the mean values as independent samples and discarding the estimated confidence intervals), and the results are reported in Table 1. All linear models have an estimated positive slope, with a significant p-value for the 3 cases: the breast collection (both cell types and subtypes) and the heart collection for cell subtypes. This confirms that the classification performance increases as the rarity of the cell types increases.

**Table 1:** Parameters of the linear regression models fitting the F1 score as a function of cell type rarity. Significant *p*-values are in bold.

| Organ | Classification level | Intercept | Slope | $p$-value of slope | Adjusted $R^2$ |
| --- | --- | --- | --- | --- | --- |
| Heart | Cell type | -0.007 | 0.131 | 0.531 | -0.047 |
| Lung | Cell type | -0.585 | 0.718 | 0.211 | 0.042 |
| Breast | Cell type | -0.104 | 0.344 | **0.013** | 0.368 |
| Heart | Cell subtype | -0.143 | 0.291 | **0.018** | 0.224 |
| Lung | Cell subtype | -0.104 | 0.194 | 0.300 | 0.006 |
| Breast | Cell subtype | -0.052 | 0.249 | **0.001** | 0.408 |

### 3.2 Swarm learning performs on par with centralized models

#### 3.2.1 Classification performance across cell types using Swarm learning

The SL classifier is compared to the LL classifier trained on 3 studies (“Local_3”). Figure 5 shows the weighted F1 score in all cell types and subtypes for LL and SL. The SL setting is directly compared to the corresponding LL setting Local_3, while Local_1 and Local_2 are also included for comparison purposes. The difference in distribution between SL and LL F1 scores is non-significant (*p*-value < 0.05) in all settings using a two-sided Mannwhitney test. When comparing Local_3 with SL in each organ, the values are 0.958 versus 0.934, and 0.972 versus 0.966 for the heart; 0.982 versus 0.982, and 0.958 versus 0.958 for the lung; 0.970 versus 0.809, and 0.796 versus 0.808 for the breast datasets, respectively. The numeric prediction accuracy values for all settings, as well as their confidence intervals, are reported in Supplementary Table 4.

**Fig. 5:**
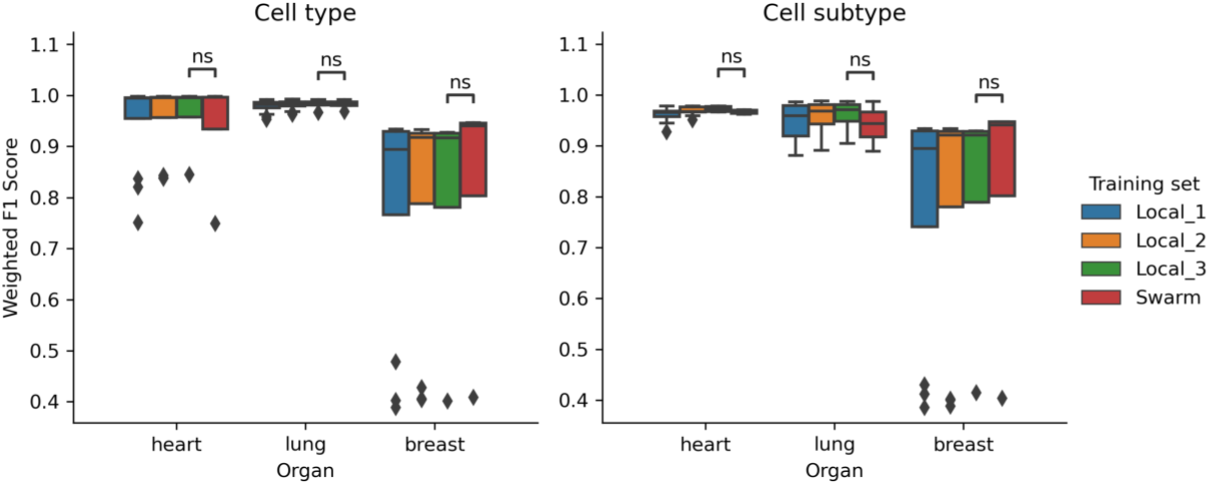
Weighted F1 score across all cell types and subtypes for LL and SL settings. Significance levels are provided by Mann-Whitney tests. Overall, the performance of local and Swarm learning approaches are comparable, showing that there is no performance loss even when training is done without sharing datasets and models in the Swarm learning setting.

#### 3.2.2 Classification performance per cell type using Swarm learning

The classification accuracy shows largely high accuracy for most cell types across the three organs as shown in the normalized confusion matrices (Figure 6 and Supplementary Figure S6). For heart, SL performs on par with LL for all cell types except Ischemic cells, Epicardium, and, to a lesser extent, Lymphatic ECs. Epicardium cells are present in low quantity and their small sample sizes (217, 0, 9, and 61) hinders efficient classification, which is reflected in lower SL performance, while “Ischemic cells” and “Lymphatic ECs” are difficult to differentiate using expression profiles. For lung, SL performs as well as LL for all cell types except respiratory basal cells, which are classified mainly as epithelial cells. This is explained by the fact that this cell type is present in small numbers (262, 245, 50, and 11, the smallest sample size for this collection). Finally, for the breast datasets, the SL classifier performs overall as well as LL. It performs on par or outperforms slightly for well-classified cell types (basal-myoepithelial cells of mammary gland, endothelial cell of lymphatic vessel, fibroblast of mammary gland, liminal hormone-sensing cell of mammary gland, macrophage, mammary gland epithelial cell, mature alpha-beta T cell, perivascular cell), it outperforms for blood vessel endothelial cells, and it slightly under-performs for some cell types which are already poorly classified by LL (capillary endothelial cell, endothelial tip cell, mature B cell, naive thymus-derived CD4-positive, alpha-beta T cell, and plasma cell). Overall, as more data are added to the collections, the “rare” cell types are better classified by LL, and thus also by SL. For LL, blood vessel endothelial cells are predicted as endothelial cells. Furthermore, several cell types are predicted as mammary gland epithelial cell: mature B cell, and two T cell types (mature alpha-beta T cell and naive thymus-derived CD4-positive, alpha-beta T cell). The very specific cell type naive thymus-derived CD4-positive, alpha-beta T cell has prediction scattered over 5 cell types. In LL setting, endothelial tip cells are predicted as mammary gland epithelial cell. For SL, in comparison, some cell types have their prediction performances significantly degraded: capillary endothelial cells, mature B cells, naive thymus-derived CD4-positive, alpha-beta T cells (which is a difficult case, even for Local), and plasma cells, which were well classified in the Local_3 setting.

**Fig. 6:**
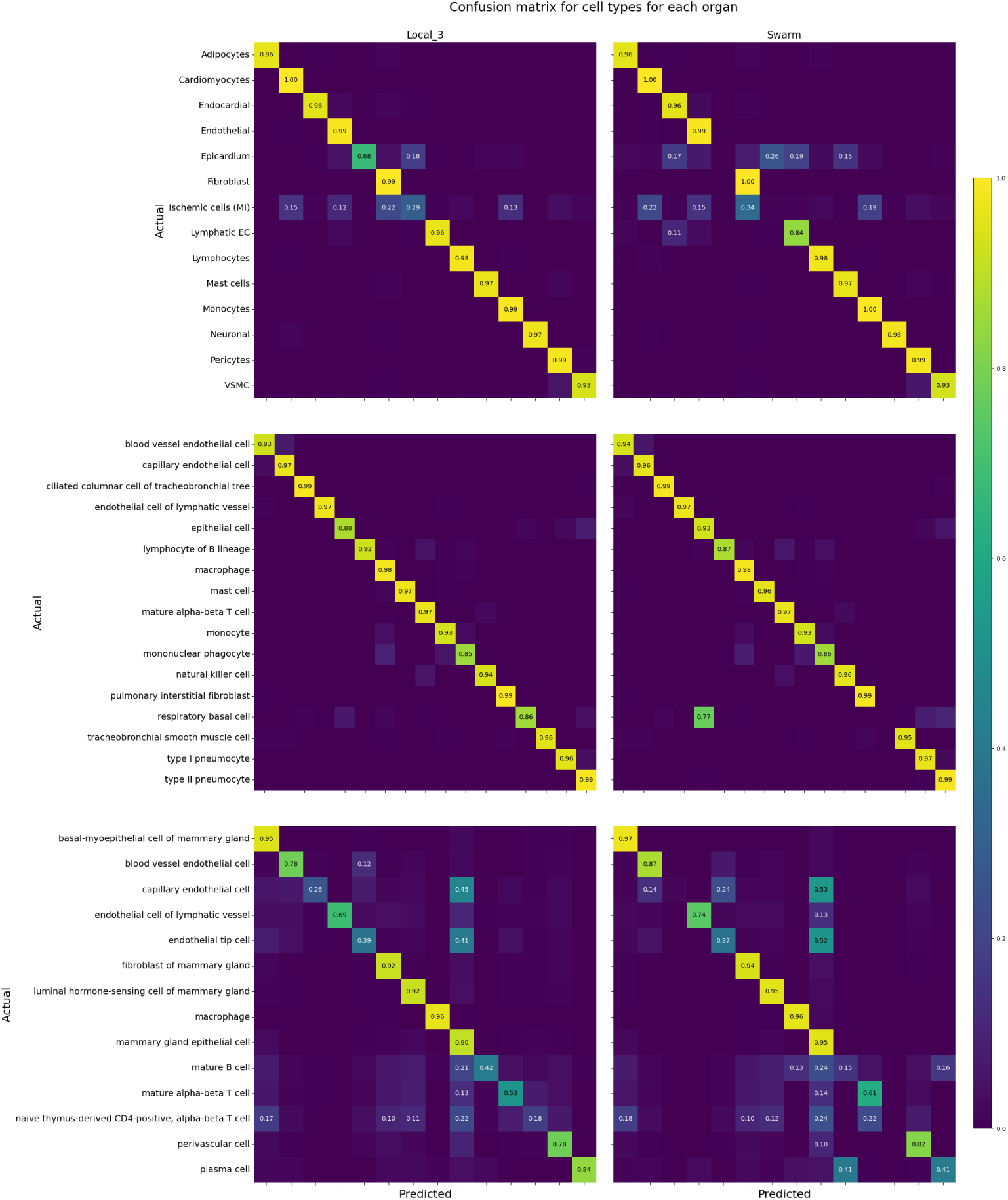
Confusion matrices for classifying cell types using Local_3 (left) versus Swarm_3 (right) classifier on heart (top), lung (center), and breast (bottom) datasets. The accuracies are are averaged over all simulation runs and normalized by row.

Visualization of the classification prediction score is available in Supplementary Figure S13.

#### 3.2.3 Cell subtype classification is challenging

Cell subtypes are defined as cell-states in a given cell lineage or main cell type that are closely related to each other and share biological properties with other cell subtypes identified within the main cell types.

For heart, in LL setting, the only cell subtype that is misclassified in the relative majority of the time is dendritic cells, which are predicted to be macrophages 47% of the time. In SL, this cell subtype is correctly classified < 10% of the time and is predicted as macrophages a (absolute) majority of the time. The other misclassified subtypes are: CD8-positive, alphabeta regulatory T cell and T cell, jointly; circulating angiogenic cell and endothelial cells, jointly; lymphocytes, as alpha-beta regulatory T cells, T cells, and natural killer cells, and plasma cells as natural kill cells, and myofibroblast cells as fibroblasts. For SL setting, 7 cell subtypes have an average accuracy < 10% (B cells, CD8-positive, alpha-beta regulatory T cells, circulating angiogenic cells, dendritic cells, lymphocytes, megakaryocytes, and plasma cells), and another subtype has an accuracy < 50% (myofibroblast cells). All of these subtypes also have low prediction accuracy in LL, with the notable addition of B cells (0.84% accuracy in LL), classified by SL mostly as monocyte and natural killer cells. All other cell subtypes with accuracy greater than 70% in LL had similar or better accuracy in SL.

In the lung datasets, in LL setting, 13% of CD1c-positive myeloid dendritic cells were predicted as lung macrophages; 16% of CD4-positive, alpha-beta T cells were predicted as and CD8-positive, alpha-beta T cells; 14% of elicited macrophages as alveolar macrophages; 12% of epithelial cell of alveolus of lung as epithelial cell of lower respiratory tract and 18% as type II pneumocytes; 15% of non-classical monocytes as classical monocytes; and 11% of pulmonary artery endothelial cells as capillary endothelial cells. In SL, the six aforementioned cell subtypes show a drop in accuracy, with epithelial cells of alveolus of lung having the largest drop in accuracy from 65% to 13%. Notably, most subtypes well classified in LL are also well classified in SL, with respiratory basal cells standing out, having an accuracy of 82% in local learning down to below 10% in Swarm learning.

In breast datasets, only seven subtypes have an accuracy over 80% in LL: basalmyoepithelial cells of mammary gland, fibroblasts of mammary gland, luminal adaptive secretory precursor cell of mammary gland, macrophages, plasma cells, and vein endothelial cells. Of the 11 subtypes which are classified with > 50% accuracy in LL, 10 are classified with an increased or equal accuracy (the outlier is mature NK T cells, which drop from an accuracy of 50% to < 10%). On the opposite side, of the 11 subtypes which are classified with < 50% accuracy in LL, 10 are classified with decreased accuracy in SL (the outlier is CD8-positive, alpha-beta memory T cells, which increase from 47% to 60%). Furthermore, for breast in LL, 6 subtypes are misclassified as luminal adaptive secretory precursor cells of mammary gland in a relative majority of cases: Tc1 cells (20%), capillary endothelial cells (41%), class switched memory B cells (25%), mammary gland epithelial cells (misclassified in an absolute majority of cases, 51%), naive thymus-derived CD4-positive, alpha-beta T cells (21%), and unswitched memory B cells (15%). This could be explained by a difficulty in finding expression signatures which differentiate well these subtypes from the biomarkers of luminal adaptive secretory precursor cells of mammary gland. These results are carried over to SL, in which the misclassification rates increase in these 6 subtypes.

Taken together, these results highlight that SL performs worse than LL when the cell subtypes are poorly classified in LL and performs better when the cell subtypes are sufficiently well classified by LL.

## 4 Discussion

Our study shows that there were no significant differences in the precision of cell type prediction between the Swarm model and the model trained on all combined data. This suggests that using the Swarm model approach is efficient method for predicting cell types in large-scale single-cell transcriptomics datasets while maintaining data privacy. However, challenges remain for predicting cell-types that are not clearly differentiable. This could be due to low sample count, lineage that is a mixture of multiple cell-types e.g. ischemic cells, overor underclustering, or closely related cell-types. Figures 6 and S6 highlight *(i)* the challenge in annotating cell types which are present in small numbers and *(ii)* the higher accuracy of annotating cell types which are well-defined and present in higher numbers.

In heart datasets tested here, cluster annotated as ischemic cells was difficult to classify in both LL and SL settings. This is due to ischemic cells being a aggregated cluster comprising of several lineages including cardiomyocytes, fibroblasts and endothelial cells. Additionally, lymphatic endothelial cells are misclassified as endocardial cells. Both celltypes are closely related and share marker genes [38]. For lung datasets, performance is good throughout classes for LL, except for “respiratory basal cells” (cell ontology id “CL:0002633”), which are classified as “epithelial cells” (cell ontology if “CL:0000066”). This is biologically sensible, as the former is a subtype of the latter in the Cell Ontology. Likewise, SL performs well across cell types except for the latter case, in which 77% of respiratory basal cells are misclassified as epithelial cells. For breast datasets, cell types including blood vessel endothelial cells, fibroblasts of mammary gland, luminal hormone-sensing cells of mammary gland, and mammary gland epithelial cells, that are well classified in LL setting, are also classified correctly in the SL setting.

The cell subtype classification task is harder than the prediction of the main cell type as different cell states can have similar marker genes (see Supplementary Figure 4). In general, the classification is fairly accurate across subtypes for the lung (1 cell subtype out of 24 with below 70% accuracy), followed by the heart (5 of 21 cell subtypes below 70% accuracy) and finally the breast (13 of 22 cell types below 70% accuracy).

This study highlights the potential of Swarm learning to build scalable and privacy-preserving models from single-cell transcriptomic data. The next step will consist in training hierarchical models for all annotation levels. Using the cell ontology as *a priori* information, hierarchical classifiers [39] can be used to annotate cells at the correct depth in the ontology. This approach will benefit from fine-grained annotated datasets which are now increasingly available to resolve automated fine-grained annotation. Another extension of this work is the building of a cross-organ model for annotation. Since many cell types are present in different organs, pooling datasets across organs will directly increase the available training size in terms of cell count per cell type, without hindering prediction of organ-specific cell types. Finally, as Swarm learning is applicable to any classifier, it can be applied to multi-omics data, leveraging different types of biological information (surface protein, chromatic accessibility, etc.) to gain a deeper insight into not only cell types, but also cell states.

## Acknowledgments

The authors gratefully acknowledge Tore Bleckwehl for providing the standardized heart atlas that was used in this study.

## Author contribution

Conceptualization: OLS, JNK, SH; Methodology: VG, KP, OLS, HK, JFZ, JNK, SH; Software: VG, KP; Formal analysis: VG, KP; Resources: JNK, RK, SH; Data Curation: VG, HK, KP; Writing - Original Draft: VG, KP, SH; Writing - Review & Editing: VG, OLS, KP, RK, JNK, SH; Visualization: VG, KP; Supervision: JNK, SH; Project administration: OLS, SH; Funding acquisition: JNK, SH;

## Supplementary Information for the paper

**Supplementary Figure S1:**
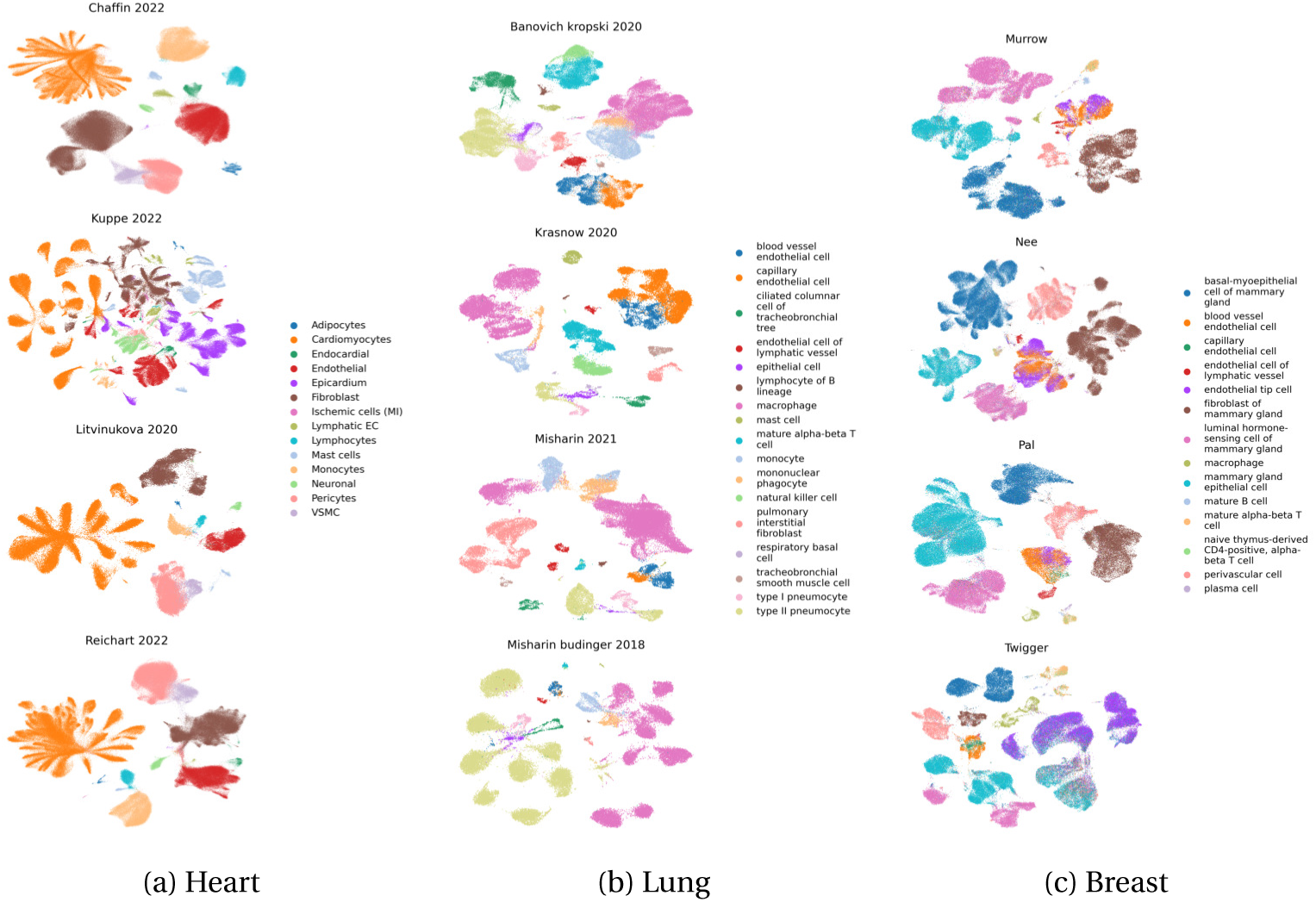
UMAPs of all datasets used in this study colored by cell type **Fig. S1:** 2D Visualization of the dataset collections, colored by the manually-curated annotation of higher level (“cell types”). UMAP projections were computed independently for each study.

**Supplementary Figure S2:**
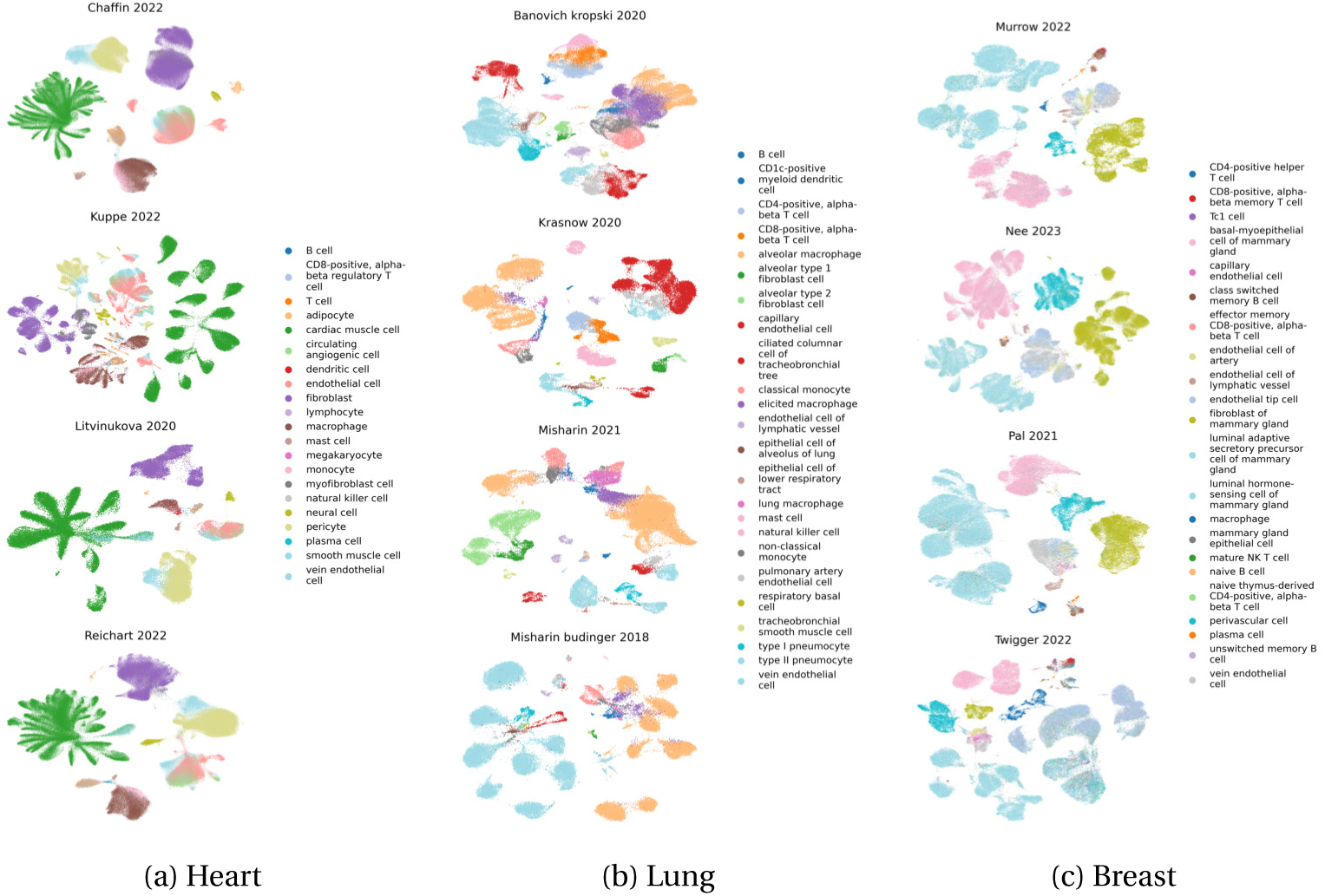
UMAP of every dataset colored by cell subtype **Fig. S2:** UMAP representation of the cell types in the heart, lung, and breast collections showing the cell subtypes.

**Supplementary Figure S3:**
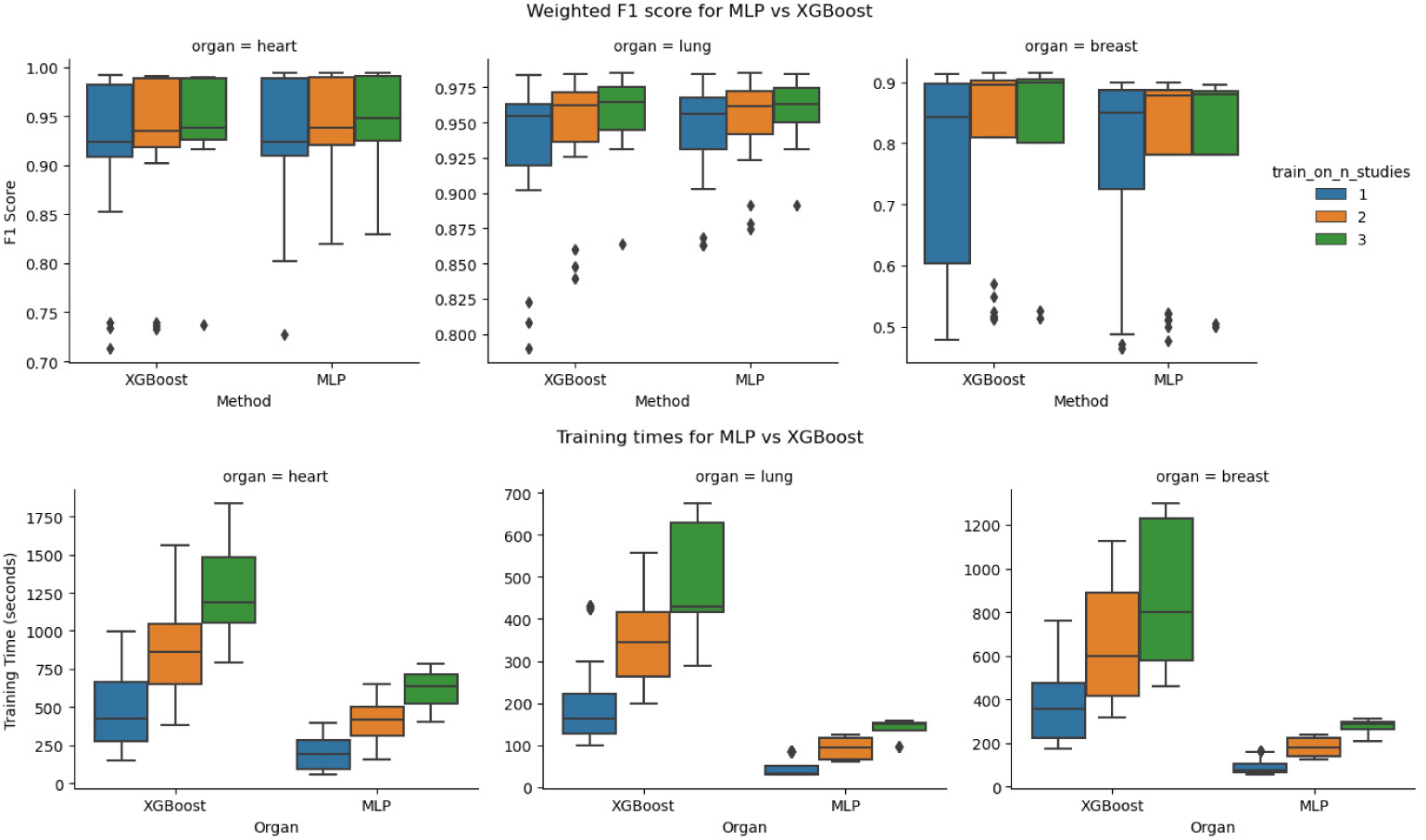
Comparison of MLP and XGBoost classifiers **Fig. S3:** Comparison of MLP and XGBoost classifiers in terms of weighted F1 score (top) and training time (bottom) at the cell type level in local learning.

**Supplementary Figure S4:**
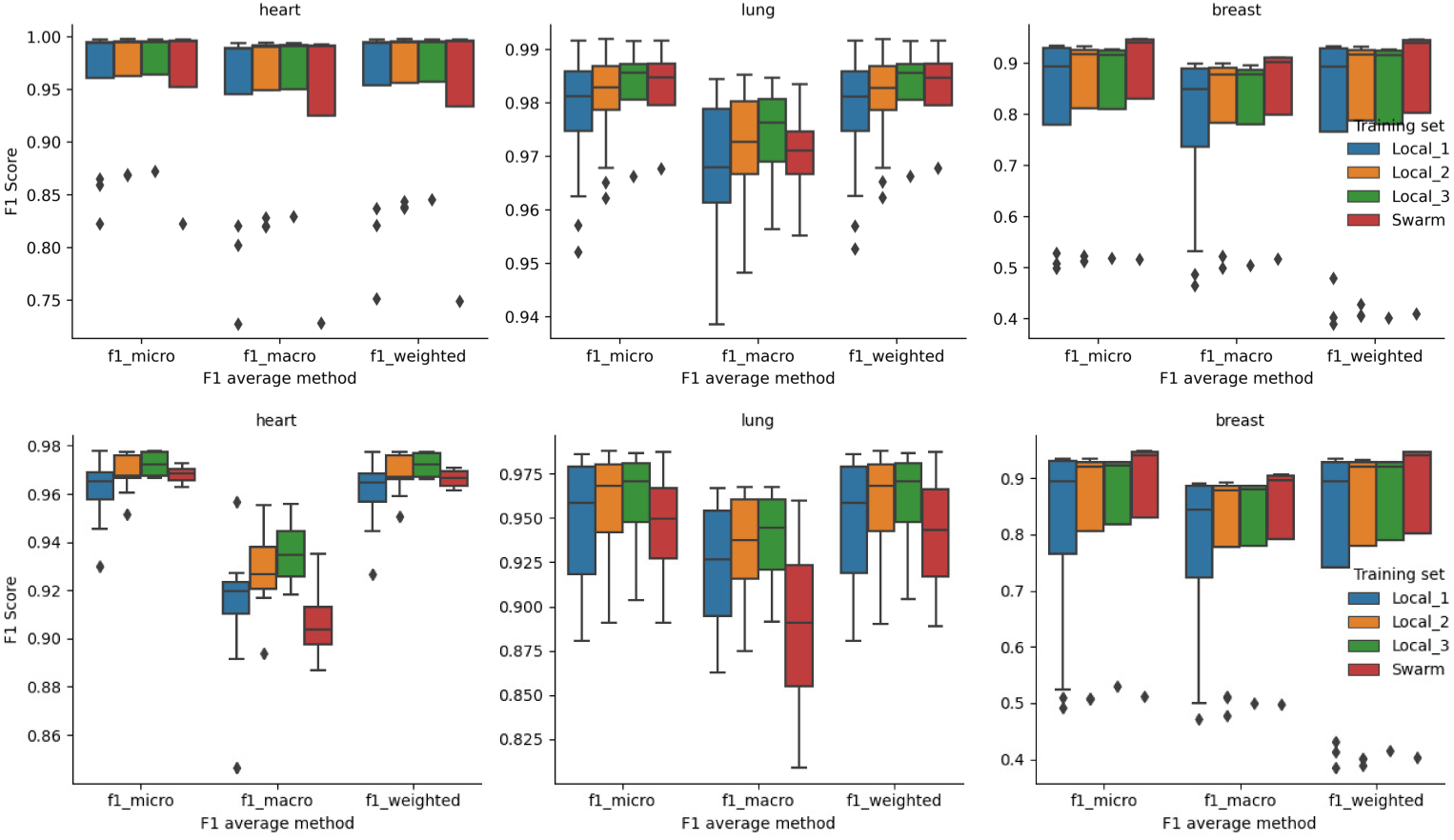
Weighted, macro, and micro F1 scores for local and swarm learning. **Fig. S4:** Micro, macro, and weighted F1 scores for cell types (top) and cell subtypes (bottom) for LL and SL. Significance levels are not shown as no Mann-Whitney *U* tests are significant.

**Supplementary Figure S5:**
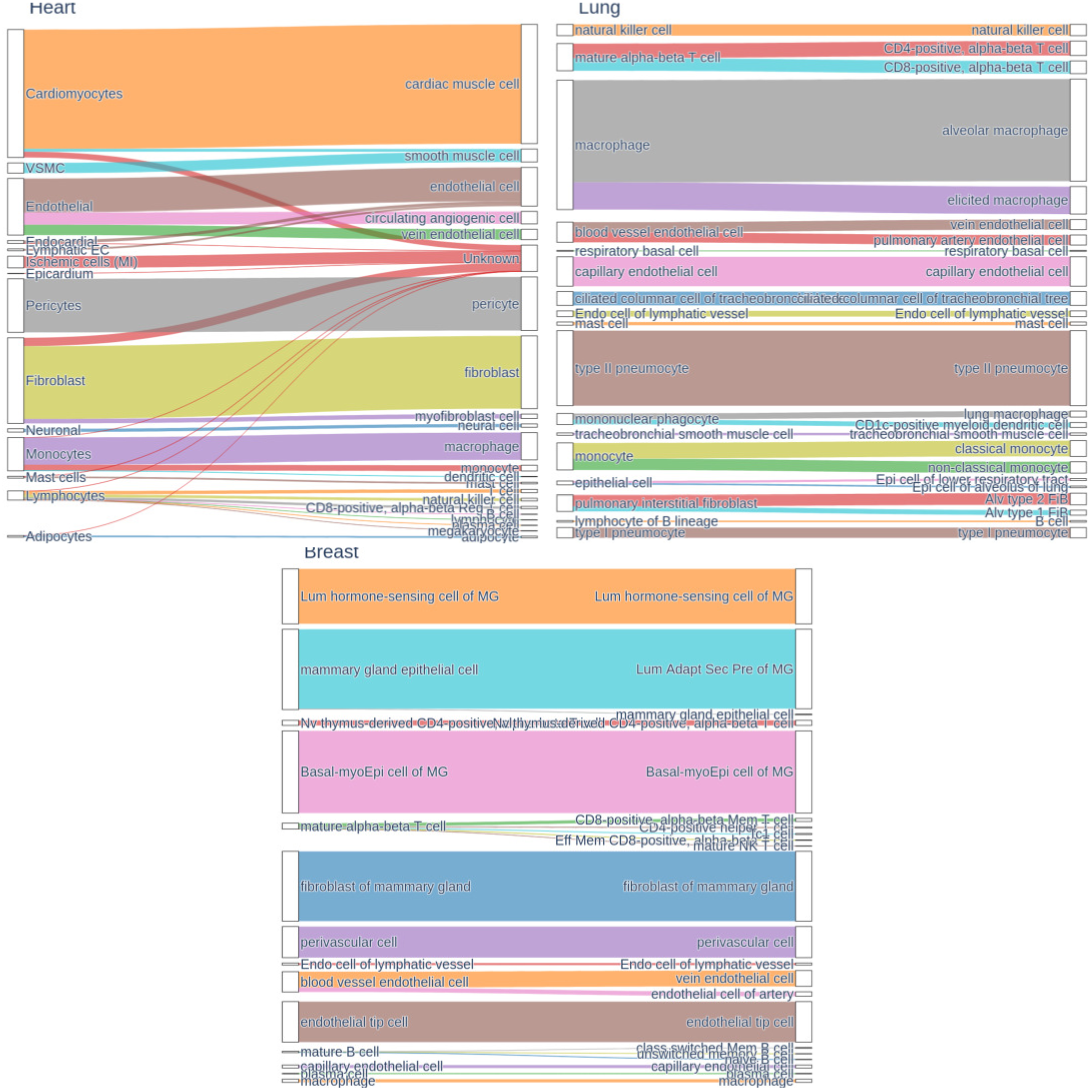
Sankey plots between cell types and subtypes **Fig. S5:** Correspondence between cell types and subtypes for each organ.

**Supplementary Figure S6:**
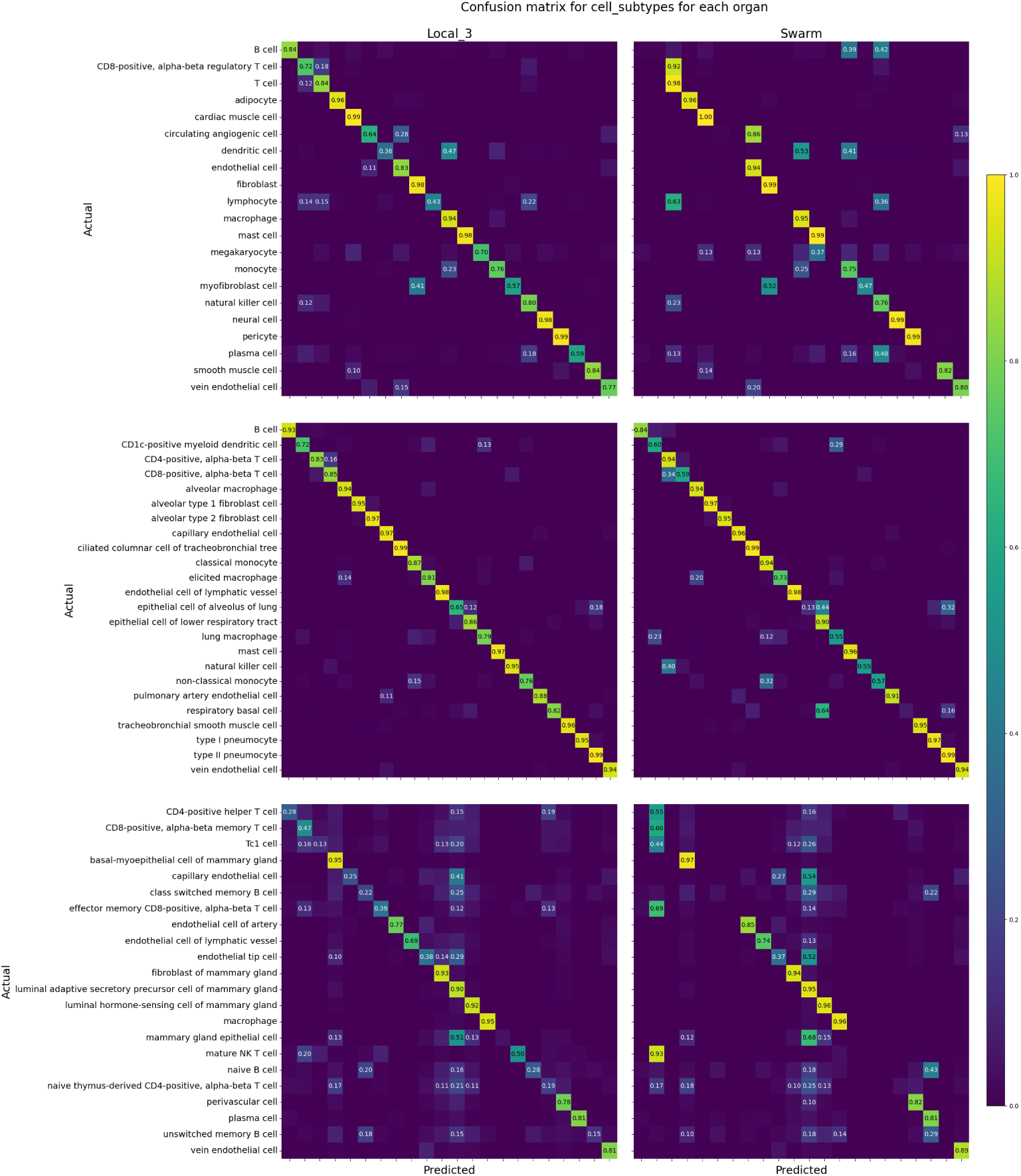
Confusion matrices for classifying cell subtypes **Fig. S6:** Confusion matrices of Local_3 (left) versus Swarm (right) at cell subtype level for the heart (top), lung (center), and breast (bottom) datasets. The accuracies are averaged over all simulation runs and are normalized by row.

**Supplementary Figure S7:**
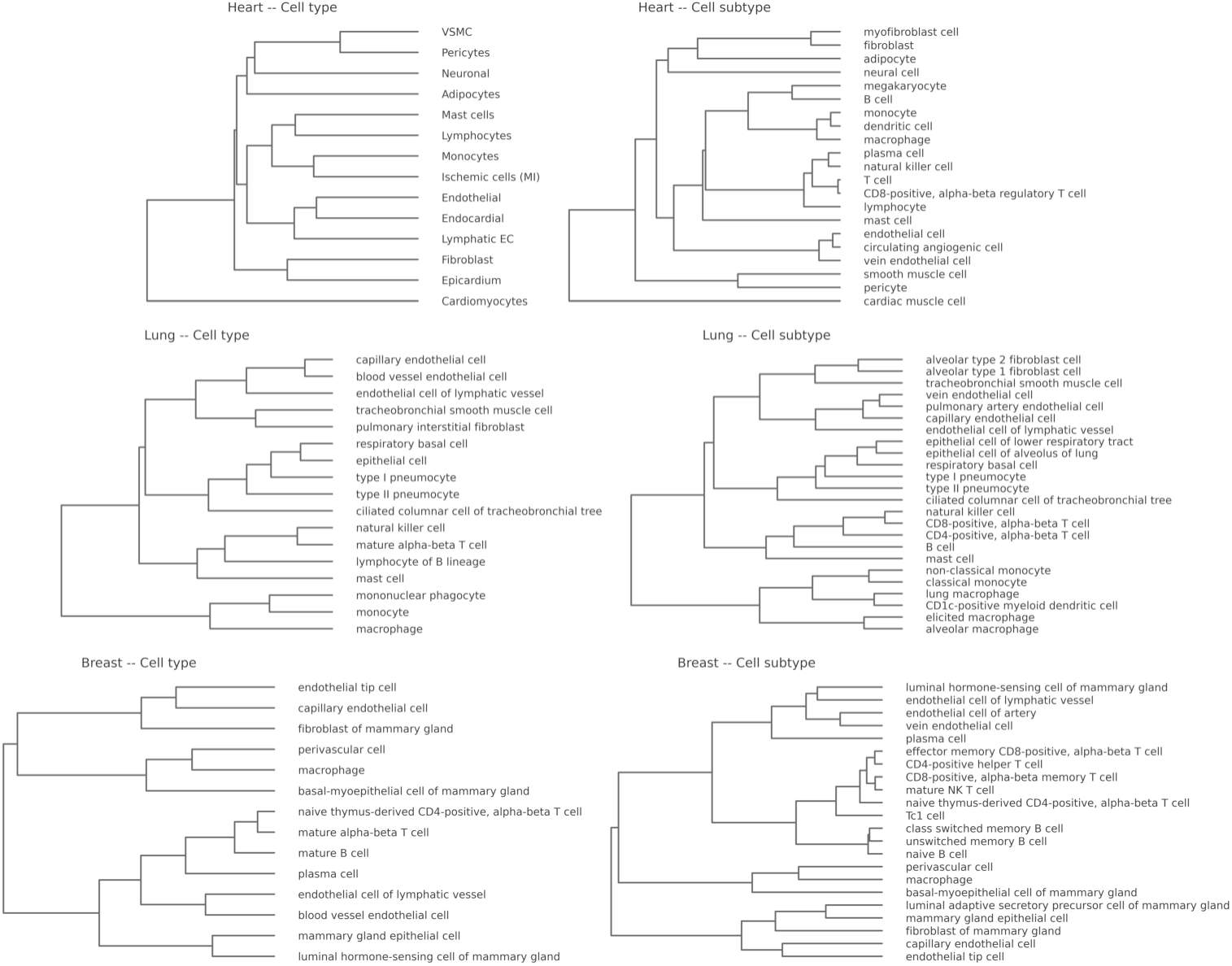
Dendrograms of cell labels **Fig. S7:** Ontology of cell types and subtypes obtained by hierarchical clustering of collections after dataset integration.

**Supplementary Figure S8:**
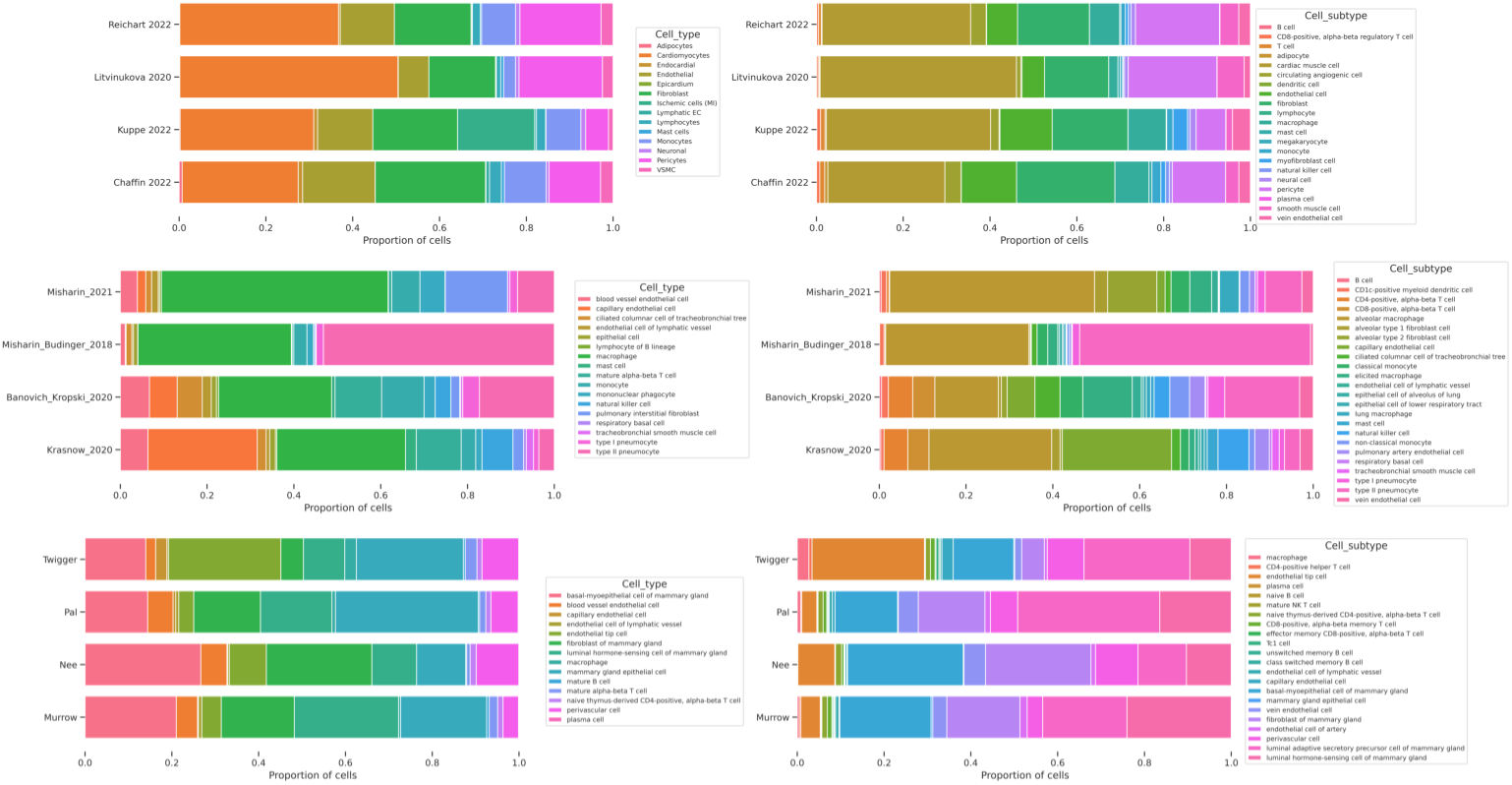
Cell type composition per dataset **Fig. S8:** Cell composition of all datasets, at both type and subtype levels.

**Supplementary Figure S9:**
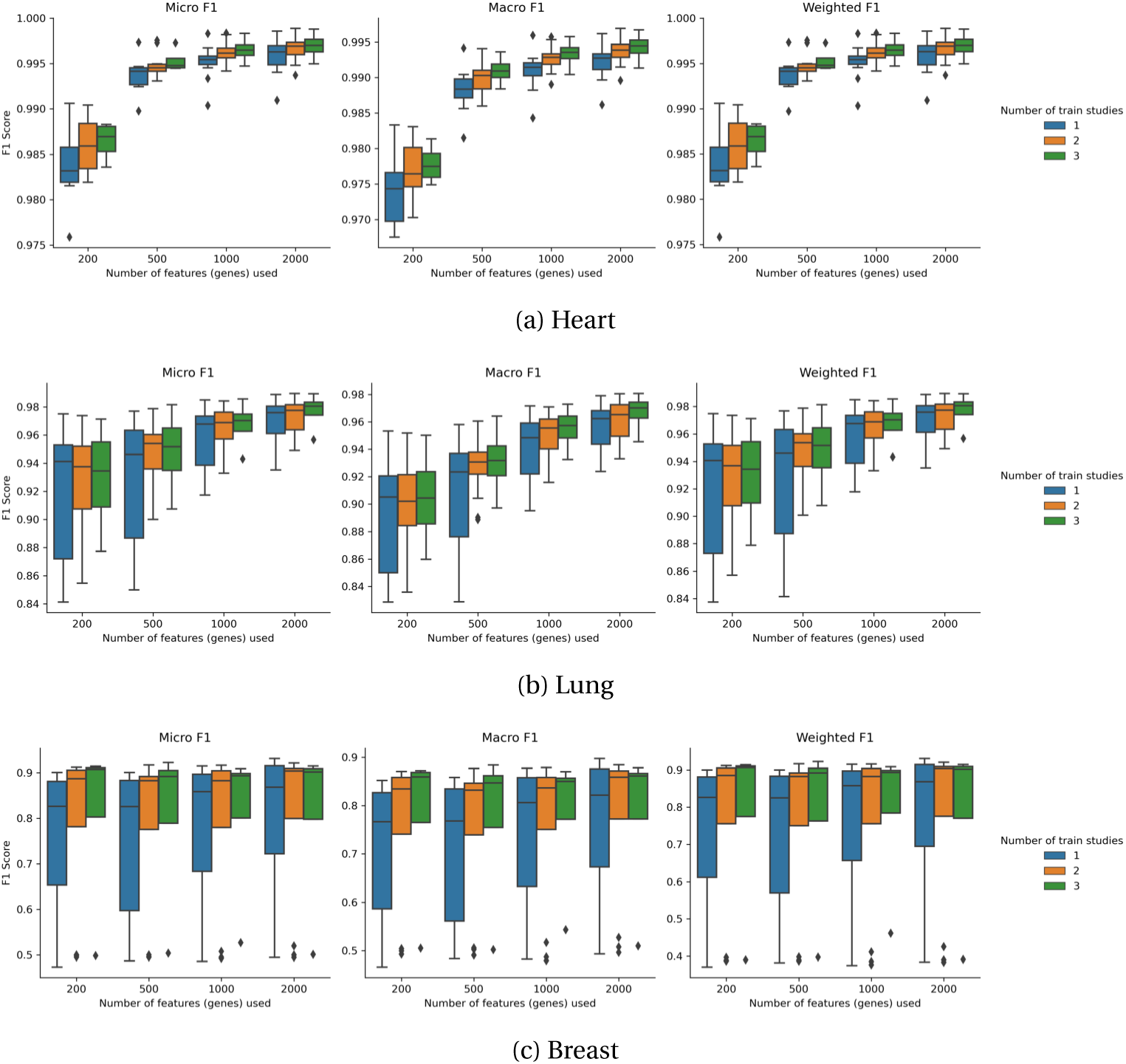
Effect of the number of HVGs **Fig. S9:** Classification performance for various numbers of HVGs selected at preprocessing.

**Supplementary Figure S10:**
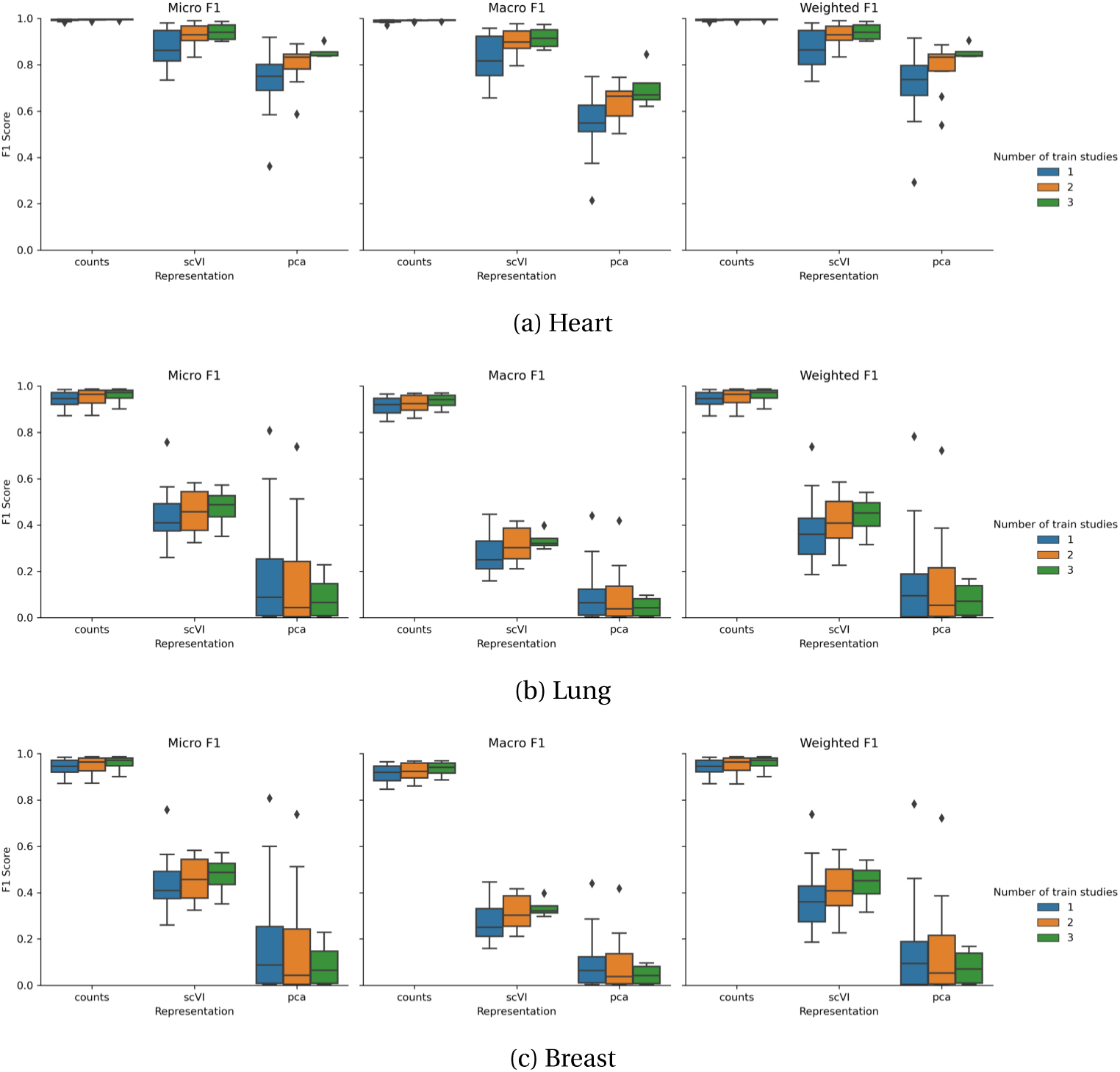
Effect of using a low-dimensional embedding Our classifier learns cell type compositions in the presence of batch effects between datasets without needing to account for these batch effects. This is possible because the feature space is the (normalized) counts. If a low-dimensional embedding were used instead, the classifier could not recover batch-effect agnostic decision boundaries. This is illustrated by the classification performance of the MLP classifier trained on (i) the normalized counts, (ii) the PCA of the counts using 50 PCs, and (iii) the embedding provided by scVI (Figure S10). The classification performance is significantly lower when using the low-dimensional embeddings, across organs and F1 scores average methods, with an even lower performance for the scVI embedding. This suggests that in the presence of batch effects, the low-dimensional embeddings do not capture the relevant information for cell type classification, and that the classifier is not able to recover the cell type composition in the presence of batch effects when using these embeddings. **Fig. S10:** Classification performance using different representations of the data.

**Supplementary Figure S11:**
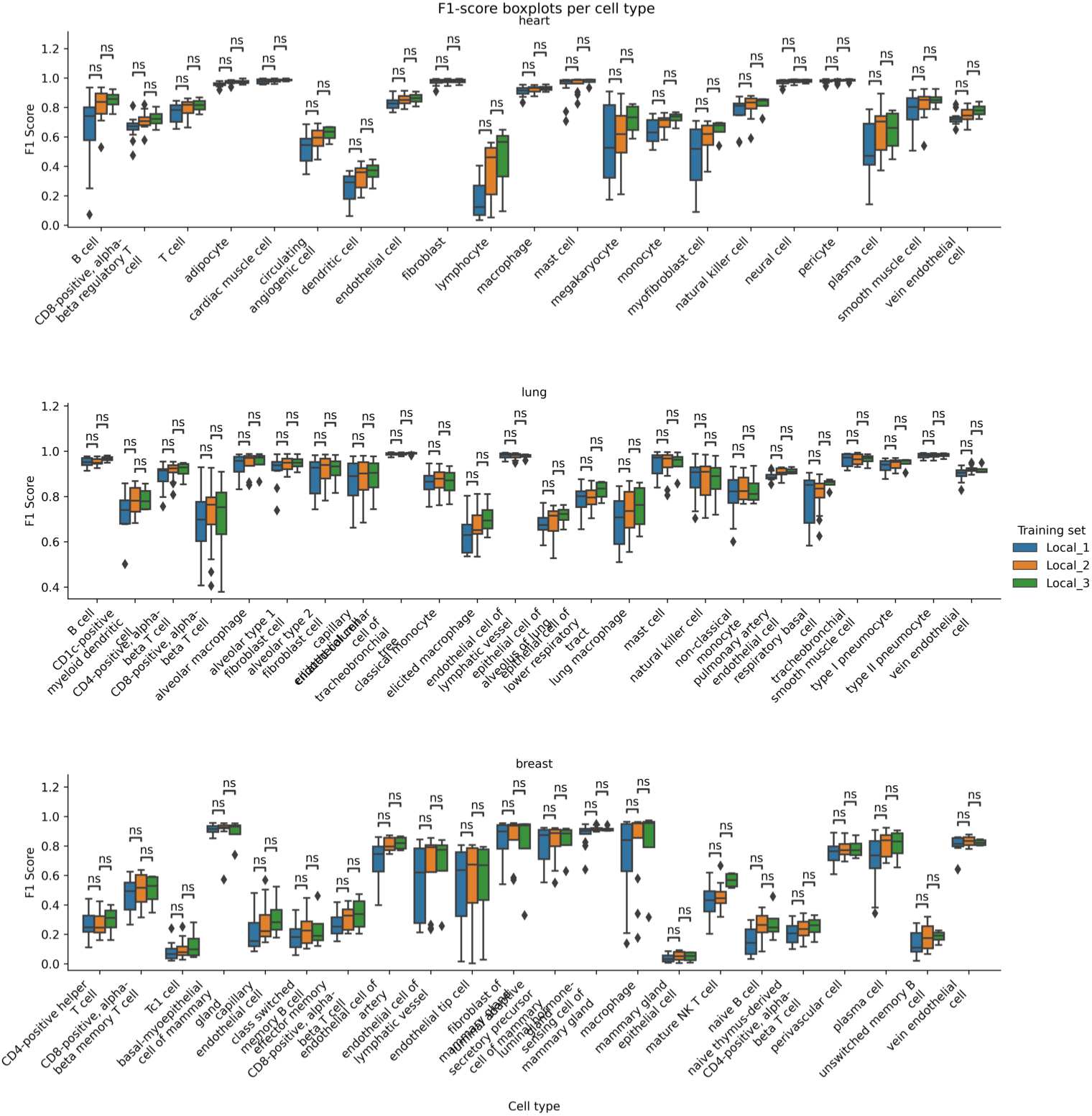
Local learning F1 scores per cell subtype **Fig. S11:** F1 score for each cell subtype with LL when training on an increasing number of studies.

**Supplementary Table 1:** Composition of datasets by cell and cell types. **Table 1**: Datasets used for training and testing SwarmMAP. Each data collection concerns one organ and is composed of 4 datasets, called “studies”. For heart, some cells had unknown subtype labels and were filtered out when using subtypes. The corresponding sample sizes are provided in parentheses.

| Collection | Organ | Study | Number of cells | Number of cell types | Number of cell subtypes |
| --- | --- | --- | --- | --- | --- |
| 1 | Heart | Chaffin 2022 [40] | 560696<br>(542216) | 14 | 21 |
|  |  | Kuppe 2022 [6] | 189349<br>(146325) | 13 | 21 |
|  |  | Litvinukova 2020 [41] | 84953<br>(79916) | 14 | 21 |
|  |  | Reichart 2022 [42] | 361649<br>(349045) | 14 | 21 |
| 2 | Lung [4] | Banovich Kropski 2020 [43] | 115249 | 17 | 24 |
|  |  | Krasnow 2020 [44] | 39901 | 17 | 24 |
|  |  | Misharin 2021 [45] | 47301 | 17 | 24 |
|  |  | Misharin Budinger 2018 [46] | 40580 | 17 | 24 |
| 3 | Breast [9] | Murrow 2022 [47] | 80726 | 14 | 22 |
|  |  | Nee 2023 [48] | 219239 | 14 | 22 |
|  |  | Pal 2021 [49] | 116432 | 14 | 22 |
|  |  | Twigger 2022 [50] | 100168 | 14 | 22 |

**Supplementary Table S12:**
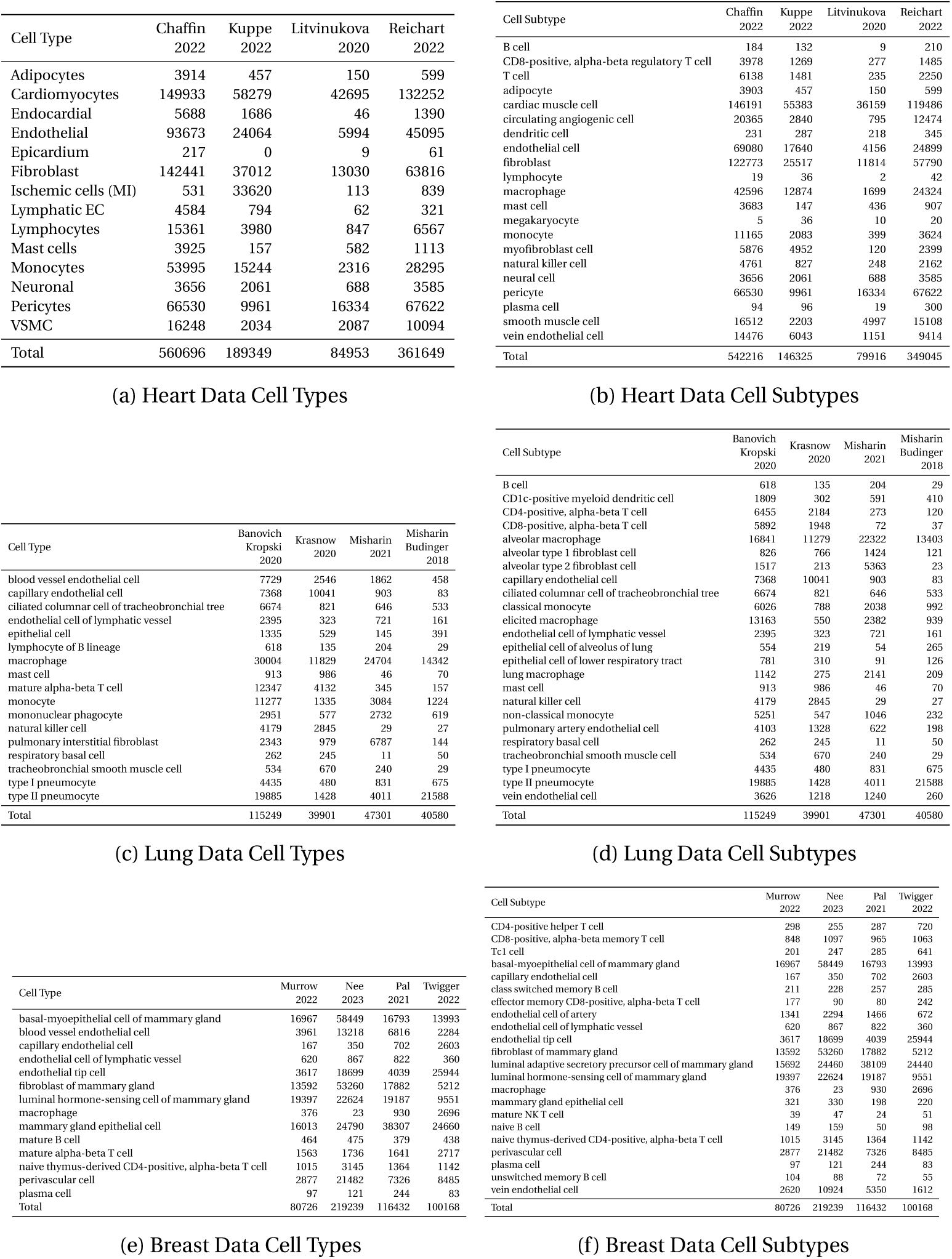
Detailed composition of datasets **Fig. S12:** Number of cell for each cell type and subtypes, for each organ.

**Supplementary Table 2:** Number of donors in each dataset. **Table 2**: Number of donors for each organ and study.

| Organ | Study | Number of donors |
| --- | --- | --- |
| Heart | Chaffin 2022 | 42 |
|  | Kuppe 2022 | 20 |
|  | Litvinukova 2020 | 14 |
|  | Reichart 2022 | 68 |
| Lung | Banovich Kropski 2020 | 38 |
|  | Krasnow 2020 | 3 |
|  | Misharin 2021 | 2 |
|  | Misharin Budinger 2018 | 8 |
| Breast | Murrow 2022 | 28 |
|  | Nee 2023 | 22 |
|  | Pal 2021 | 21 |
|  | Twigger 2022 | 18 |
| Total |  | 284 |

**Supplementary Table 3:** Data download links. **Table 3**: List of human heart, lung, and breast datasets used in this study. Download links of all datasets for CellxGene platform are provided.

| Collection | Organ | Dataset | Download link |
| --- | --- | --- | --- |
| 1 | Heart | Chaffin 2022 | https://singlecell.broadinstitute.org/single_cell/study/SCP1303/ |
|  |  | Kuppe 2022 | https://cellxgene.cziscience.com/collections/8191c283-0816-424b-9b61-c3e1d6258a77 |
|  |  | Litvinukova 2020 | https://cellxgene.cziscience.com/collections/b52eb423-5d0d-4645-b217-e1c6d38b2e72v |
|  |  | Reichart 2022 | https://cellxgene.cziscience.com/collections/b52eb423-5d0d-4645-b217-e1c6d38b2e72e75342a8-0f3b-4ec5-8ee1-245a23e0f7cb |
| 2 | Lung | Banovich Kropski 2020 | https://cellxgene.cziscience.com/collections/b52eb423-5d0d-4645-b217-e1c6d38b2e726f6d381a-7701-4781-935c-db10d30de293 |
|  |  | Krasnow 2020 |  |
|  |  | Misharin 2021 |  |
|  |  | Misharin and Budinger 2018 |  |
| 3 | Breast | Murrow 2022 | https://cellxgene.cziscience.com/collections/b52eb423-5d0d-4645-b217-e1c6d38b2e7248259aa8-f168-4bf5-b797-af8e88da6637 |
|  |  | Nee 2023 |  |
|  |  | Pal 2021 |  |
|  |  | Twigger 2022 |  |

**Supplementary Table 4:** Averages of weighted F1 scores for LL and SL. **Table 4**: Averages of weighted F1 scores for LL and SL across all settings. Confidence intervals are computed using the t-distribution. For each setting, the highest value is highlighted in bold.

| Organ | Label | Training set | Weighted F1 score (Confidence Interval) |
| --- | --- | --- | --- |
| Heart | Cell type | Local_1 | 0.947 (0.896, 0.997) |
|  |  | Local_2 | 0.957 (0.917, 0.996) |
|  |  | Local_3 | **0.958** (0.884, 1.032) |
|  |  | Swarm | 0.934 (0.813, 1.056) |
| Heart | Cell subtype | Local_1 | 0.961 (0.953, 0.968) |
|  |  | Local_2 | 0.968 (0.964, 0.973) |
|  |  | Local_3 | **0.972** (0.966, 0.978) |
|  |  | Swarm | 0.966 (0.962, 0.971) |
| Lung | Cell type | Local_1 | 0.978 (0.97, 0.985) |
|  |  | Local_2 | 0.981 (0.975, 0.987) |
|  |  | Local_3 | **0.982** (0.971, 0.993) |
|  |  | Swarm | **0.982** (0.972, 0.992) |
| Lung | Cell subtype | Local_1 | 0.945 (0.922, 0.969) |
|  |  | Local_2 | 0.954 (0.933, 0.975) |
|  |  | Local_3 | **0.958** (0.921, 0.995) |
|  |  | Swarm | 0.941 (0.899, 0.982) |
| Breast | Cell type | Local_1 | 0.786 (0.661, 0.91) |
|  |  | Local_2 | 0.794 (0.664, 0.924) |
|  |  | Local_3 | 0.79 (0.536, 1.044) |
|  |  | Swarm | **0.809** (0.547, 1.07) |
| Breast | Cell subtype | Local_1 | 0.78 (0.653, 0.908) |
|  |  | Local_2 | 0.791 (0.657, 0.926) |
|  |  | Local_3 | 0.796 (0.547, 1.046) |
|  |  | Swarm | **0.808** (0.544, 1.072) |

**Supplementary Figure S13:**
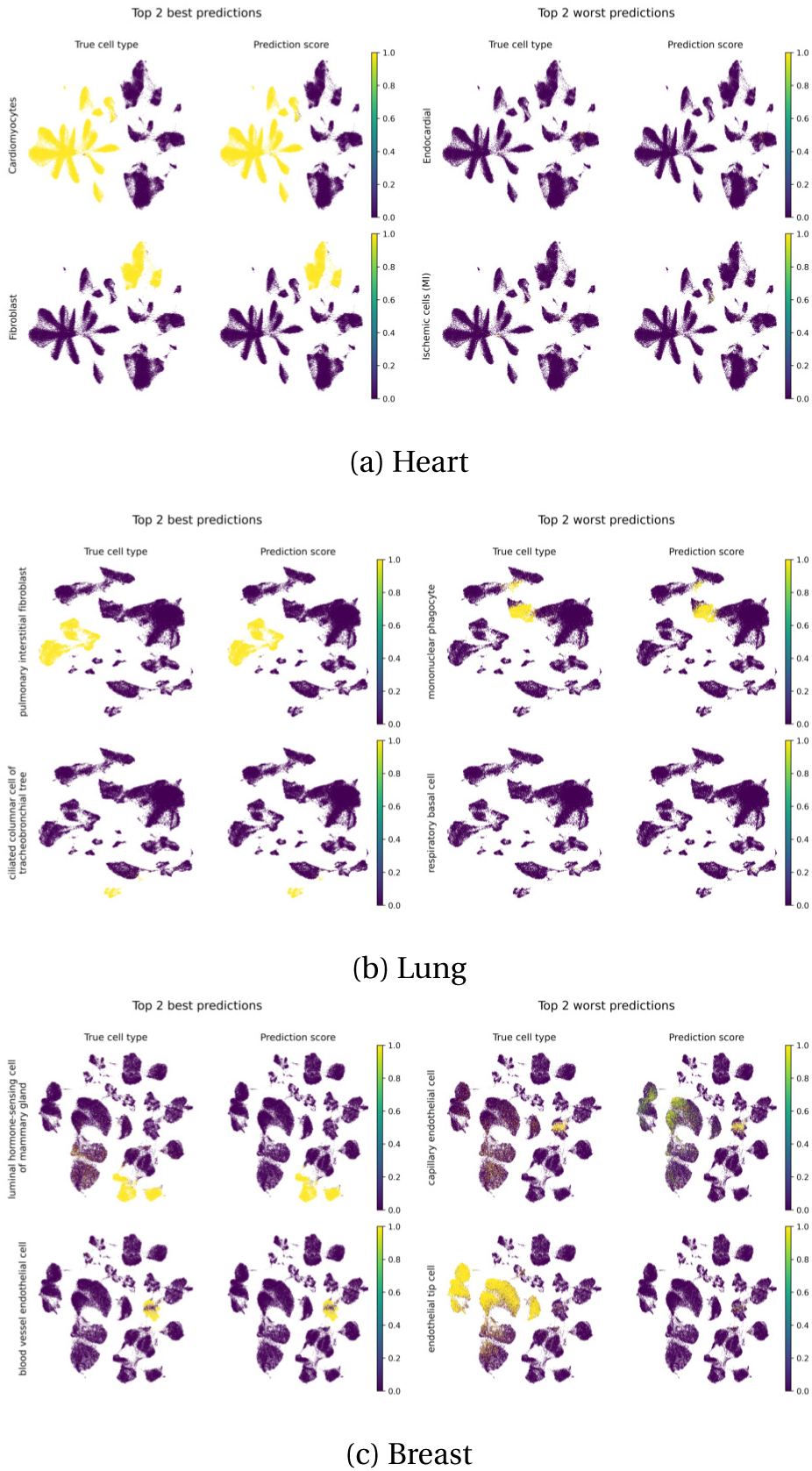
Visualizing classification prediction score Both local and Swarm classifiers display heterogeneous classification performance across cell types. To understand the reasons thereof, Figure S13 compares the true label with its prediction score. The results are represented for the two best classified and two worst classified cell types, for every organ. The values are represented in the UMAP of the test study. On one experiment experiment from Local_3 is considered for each organ and the test studies used are “Litvinukova 2020”, “Misharin 2021”, and “Twigger 2022” for heart, lung, and breast collections, respectively. **Fig. S13:** Top 2 best (left) and worst (right) classified cell types for lung. For each row, the left pane shows the true cell types and the right pane shows the prediction score.

## Footnotes

1 https://cellxgene.cziscience.com/datasets

1 https://singlecell.broadinstitute.org/single_cell

